# The completeness and stratification in yeast genotype-phenotype space

**DOI:** 10.1101/2020.01.13.904607

**Authors:** Jianguo Wang, Xionglei He

**Affiliations:** State Key Laboratory of Biocontrol, School of Life Sciences, Sun Yat-sen University, Guangzhou 510275, China

## Abstract

Genotype and phenotype are two themes of modern biology. While the running principles in genotype has been well understood (e.g., DNA double helix structure, genetic code, central dogma, etc.), much less is known about the rules in phenotype. In this study we examine a yeast phenotype space that is represented by 405 quantitative traits. We show that the space is convergent with limited latent dimensions, which form surprisingly long-distance chains such that all traits are interconnected with each other. As a consequence, statistically uncorrelated traits are linearly dependent in the multi-dimensional phenotype space and can be precisely inferred from each other. Meanwhile, the performance is much poorer for similar trait inferences but from the genotype space (including DNA and mRNA), highlighting the dimension stratification between genotype space and phenotype space. Since the world we’re living is primarily phenotypic and what we truly care is phenotype, these findings call for phenotype-centered biology as a complement for the cross-space genetic thinking in current biology.

## Introduction

The physical world is both macroscopic and microscopic, the former of which is the manifestation of the latter (Fig. 1a). Physicists adopt two rather parallel frameworks to describe the world: classical mechanics for the macroscopic layer and quantum mechanics for the microscopic layer. Unification of the two frameworks became possible only after each independently became mature(*1*). For biologists, the macroscopic layer is the phenotype of an organism and the microscopic layer is the genotype. The mainstream of current biology adopts a bottom-up thinking: because genotype is the basis of phenotype, we rely on the former to understand the latter(*2*). During the past decades a quite mature discipline - molecular biology – has been developed for describing genotype and its direct products at molecular levels. However, efforts to apply genotype understandings to phenotype, the theme of modern genetics, appear to be successful only for rather simple traits(*3*). Dishearteningly, the community of genetics does not seem optimistic with the future of studying complex traits(*4-6*). Hence, a possible solution to the situation is, like what the physicists used to do, to develop a rather independent discipline that deals with only the macroscopic layer (i.e, phenotype)(*7, 8*).

**Fig. 1.**
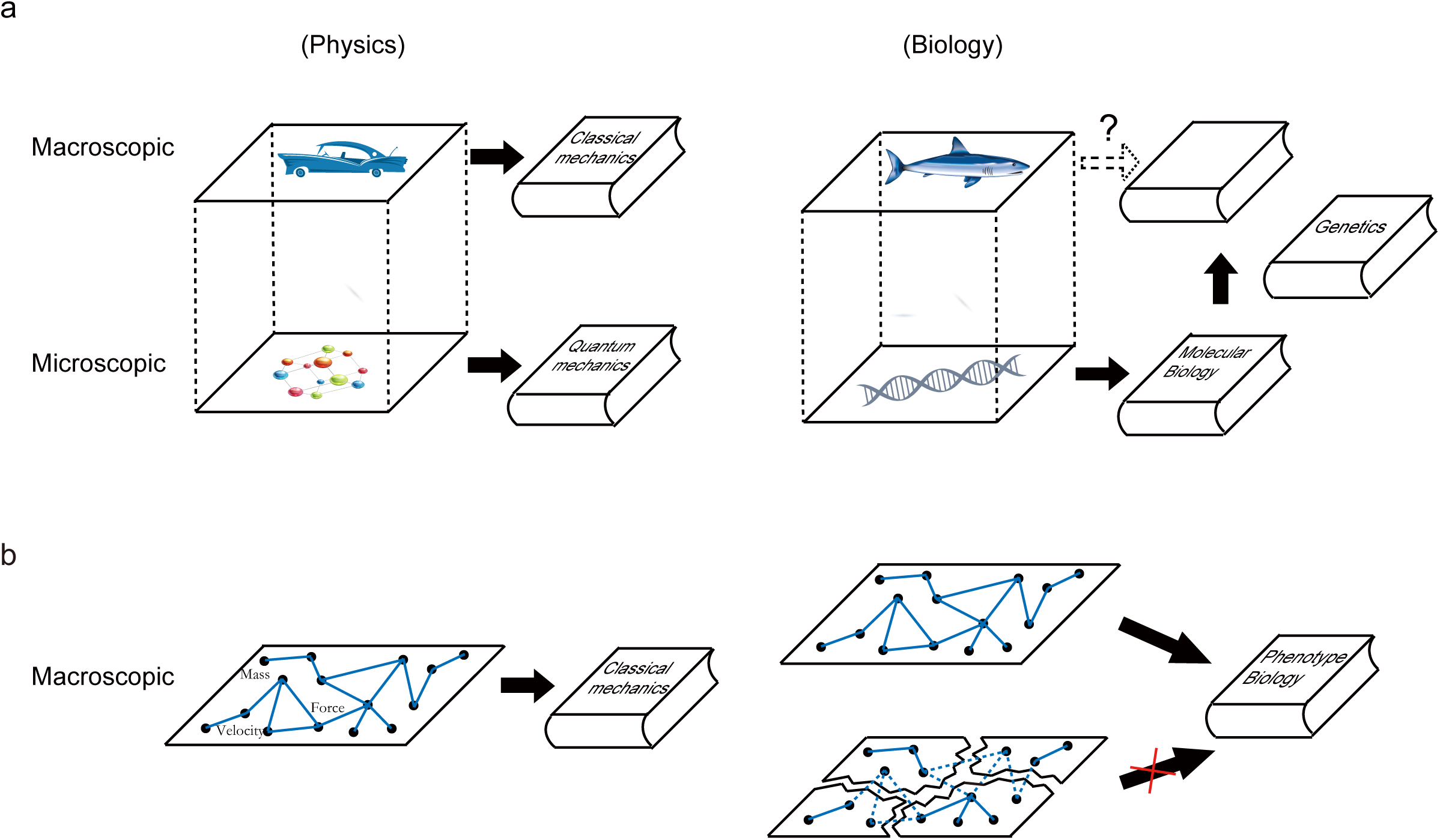
The rationale for developing phenotype biology. **(a)** Different philosophies between physics and biology in dealing with the macroscopic and microscopic layers of the examined world. **(b)** An independent discipline lies in the completeness of the logic chains in the examined space such that the various objects examined are logically interconnected rather than isolated from each other.

Available studies on phenotype space focused primarily on either the evolution of a few traits or the interaction of related traits(*9-13*). In both scenarios what people could have discovered are local logics within a phenotype space. It is unclear whether a typical phenotype space is splitting with isolated sets of local logics or is fully interconnected via global logics (Fig. 1b). The answer to this question is important, because development of phenotype biology lies in the completeness of phenotypic logics such that seemingly unrelated traits can be deterministically deduced from each other without the knowledge outside the phenotype space.

## Results

To address the question, dense sampling of a phenotype space is required. In this study we used the budding yeast *Saccharomyces cerevisiae* as a model system. Because the yeast cell morphology reflects various cellular events, including progression through the cell cycle, establishment of cell polarity, and regulation of cell size, a total of 405 traits related to cell morphology are used to represent the yeast phenotype space(*14*). It was known that at least 2,700 distinct non-essential genes are involved in regulating these traits(*14*), underscoring the cellular complexity behind this phenotype space (Fig. S1a). The 405 traits are typically about area, distance, and angle obtained based on several dozens of basic coordinate points or lines that describe the shape of mother cell and bud, the neck separating mother cell from bud, and the shape and localization of the nuclei in mother cell and bud (Fig. 2a). Because cell morphology changes with cell cycle progression, three stages roughly corresponding to G1-S phase, S-G2 phase, and cytokinesis, respectively, are considered separately. In addition, because the number of cells belonging to a stage is proportional to the duration of the stage, the frequency of cells belonging to a specific stage is also considered. Hence, the yeast phenotype space examined here comprises both cell morphology and physiology information, representing a rather comprehensive phenotypic characterization of the organism.

**Fig. 2.**
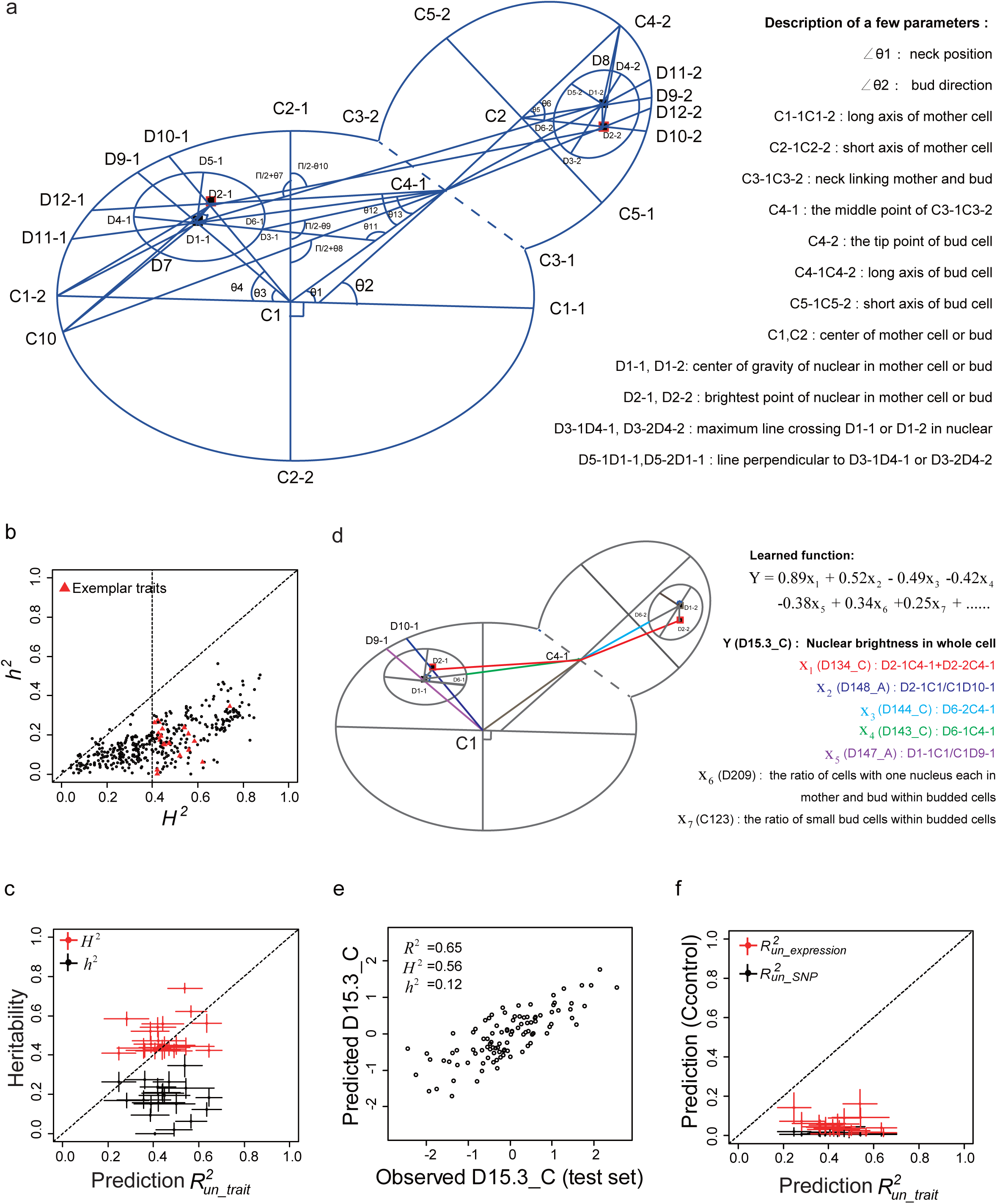
Modelling yeast morphological traits using their uncorrelated traits. **(a)** The yeast cell morphology outlined by dozens of coordinate points, lines and angles from which a total of 405 quantitative traits are derived. **(b)** The narrow-sense (*h*^2^) and broad-sense (*H*^2^) heritability of the 405 yeast traits. Twenty exemplar traits are selected from those with *H*^2^ > 0.4. **(c)** The prediction performance 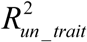 is invariably larger than *h*^2^ and comparable to *H*^2^ for each of the 20 exemplar traits. Each dot represents a trait. 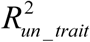 is the trait variance explained by the learned linear function in which its uncorrelated traits are explanatory variables. Error bars of *H* ^2^ and *h*^2^ are standard error estimated by Jackknife. Error bars of 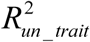 are standard deviation estimated by 100 repeats of the learning. **(d)** The learned linear function for exemplar trait D15.3_C, with details of its top seven uncorrelated explanatory traits (X_1,_ X_2_, …, X_7_) provided. **(e)** The observed and predicted trait values of D15.3_C. Data of one random set of testing segregants (*n* = 100) are shown, with each dot representing a segregant. *R*^2^ shows the square of Pearson correlation coefficient, and *H*^2^ and *h*^2^ show the broad-sense and narrow-sense heritability, respectively. **(f)** 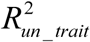 is much larger than both the uncorrelated-SNP-based prediction performance 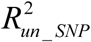 and the uncorrelated-expression-based prediction performance 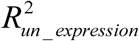 for each of the 20 exemplar traits. Each dot represents a trait. To make sure the three modeling processes are comparable, the same number of uncorrelated traits, uncorrelated SNPs, or uncorrelated gene expressions are used in the modellings for each exemplar trait. Error bars show standard deviation.

We examined a population of 815 haploid segregants that are produced by a hybrid of two *S. cerevisiae* strains(*15*). In the population the median size of linkage blocks is ∼106 kilo-base pairs, and the average genetic distance between two segregants is ∼0.25%, with a largely normal frequency distribution (Fig. S1b). Hence, the genetic heterogeneity of the population is reasonably high and the genotypes are well-mixed with no apparent structure. We obtained the data of the 405 morphology related traits of the segregants that grow in the rich medium YPD at 30 centi-degree(*16*). The traits are all quantitative, and in nearly all cases the trait values show a bell-shape density distribution (Fig. S2), reflecting the complexity of their underlying genetic architecture. We computed the broad-sense heritability (*H* ^2^) for each of the traits in the population, which measures the theoretical upper-bound of understanding phenotype from genotype. The *H*^2^ ranged from 0 to 0.86, with a median of 0.42 (Fig. 2b), suggesting that we could possibly model typically ∼42% of trait variance in the population using genotype. However, modeling broad-sense trait heritability is notoriously difficult because of the enormous complexity of gene-gene interactions. Hence, often a more realistic goal in current genetics is narrow-sense heritability (*h*^2^), which is the additive component of *H* ^2^. With genome information of the segregants we calculated *h*^2^ for each of the traits and found them generally much smaller than *H*^2^, ranging from 0 to 0.56 with a median of 0.15 (Fig. 2b).

Since many of the yeast traits are related, we first focused on 20 exemplar traits each with an *H*^2^ > 0.4 (Fig. 2b; Methods). We asked whether the 20 traits each can be modeled using their statistically uncorrelated traits. The answer would be negative if the phenotype space is splitting with only short-distance logic chains that account for apparently correlated traits. Alternatively, the phenotype space could be fully interconnected via long-distance logic chains such that statistically uncorrelated traits are deeply dependent in the high-dimensional space. For each trait we computed its Pearson’s correlation in the segregant population with the rest 404 traits, respectively. Two traits were called statistically uncorrelated if the Pearson’s *R*^*2*^ < 0.02, which corresponds to *P* > 0.01 after Bonferroni correction for multiple testing (*n* = 404). Using jackknife and bootstrapping methods we confirmed that the assignments of uncorrelated traits in the population are robust (Fig. S3-S4). We obtained on average ∼200 uncorrelated traits for each trait (Fig. S5a). We divided the population into a training subset with 715 segregants and a testing subset with 100 segregants. We learned a linear function to model an exemplar trait using its uncorrelated traits, and tested the performance of the learned function in the testing subset (Methods). Remarkably, for all exemplar traits the variances explained by the learned models are no smaller than *h*^2^, and actually in most cases are close to or even slightly higher than *H*^2^ (Fig 2c). There are often quite a few explanatory variables (i.e, uncorrelated traits with a non-zero coefficient) in each of the linear functions (Fig. S5b); the absolute coefficients are often small (Fig. S6), with no apparent correlation with the absolute Pearson’s *R*s between the explanatory traits and a focal exemplar trait (Fig. S7). A close examination of the exemplar traits with their top explanatory traits reveals no obvious interactions between them. For example, there are seven explanatory traits each with an absolute coefficient large than 0.2 in the case of D15.3_C, an exemplar trait that measures the total brightness of the two nuclei (Fig. 2d-e, Fig. S8): the seven top explanatory traits are about the relative position of the two nuclei to each other and to the center of the mother cell, as well as some cell population ratio parameters. Hence, the relationship between explanatory traits and a focal exemplar trait has to be understood with a rather complex linear function. The success of the modelling suggests statistically uncorrelated traits are actually interconnected via hidden logics in the phenotype space.

As a control, we modelled each exemplar trait in the same way but using its uncorrelated genetic loci (Single nucleotide polymorphisms or SNPs). For each of the 20 exemplar traits we identified its quantitative trait loci (QTLs). In all the cases the identified QTLs explain most of the corresponding *h*^2^ (Fig. S9a). We excluded the corresponding QTLs and used the remaining genotype (i.e., uncorrelated SNPs) to model each of the traits (Methods). We found that the trait variance explained by uncorrelated SNPs is invariably much smaller than *H* ^2^, roughly reflecting the missing portion of *h*^2^ (Fig. 2f and Fig. S9b-d). This finding is important, because it indicates that the previous successes of uncorrelated traits are not due to the lack of heterogeneity of the yeast population. It also suggests no effective logics for connecting a trait with its uncorrelated SNPs.

Furthermore, we modelled the exemplar traits using their uncorrelated gene expressions. This can be done because of recently available RNA-seq data of the yeast segregant population(*17*). For each of the ∼6,000 yeast genes there is typically a bell-shape density distribution of expression level in the population (Fig. S10). Similar to the definition of uncorrelated traits, the uncorrelated gene expressions for each exemplar trait are identified if the *P-*value of an expression-trait correlation is larger than 0.01 after Bonferroni correction for multiple testing (*n* = 5719). Then, a linear function of the uncorrelated gene expressions is learned for each of the exemplar traits (Methods). We found that the modeling performance is just comparable to that of uncorrelated SNPs, much poorer than that of uncorrelated traits (Fig. 2f and Fig. S11a). Notably, for a typical exemplar trait the number of uncorrelated gene expressions is over ten times higher than the number of uncorrelated traits (Fig. S11b); also, the cutoff *R*^2^ (=0.028) in defining uncorrelated gene expressions is slightly higher than the cutoff *R*^2^ (=0.02) in uncorrelated traits (Fig. S11c), due to the different numbers (5719 v.s. 404) used in multiple testing correction when the same corrected *P*-value cutoff (= 0.01) was used. Hence, the poorer performance of uncorrelated gene expressions relative to uncorrelated traits cannot be explained by any technical issues. Instead, it suggests a logic stratification between trait space and expression space.

The uncorrelated-trait-based modeling can be understood from a geometric point of view, in which a trait is represented by a vector and two traits are uncorrelated if the two vectors are nearly perpendicular. For simplicity we may consider three non-parallel vectors *α, β*, and *η* in a three-dimensional space, with *θ*_*1*_ representing the angle between α and *η* and *θ*_*2*_ the angle between *β* and *η* (Fig. 3a). The correlation (*R*) of two vectors, say, vector *α* and *η*, can be written as:

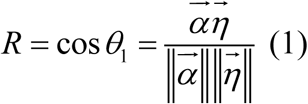

**Fig. 3.**
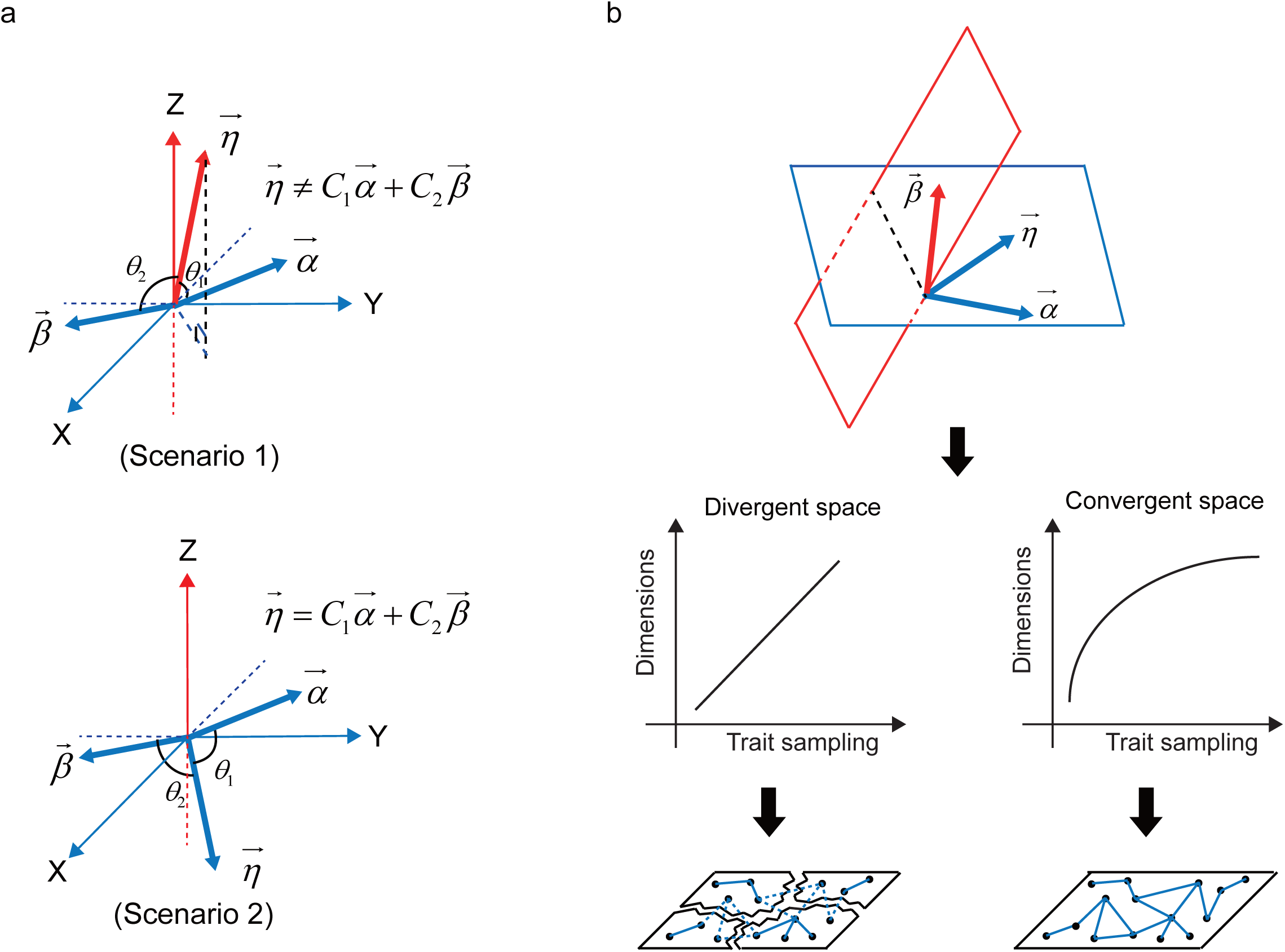
The theory for explaining linear dependence of uncorrelated traits. **(a)** Here a trait is expressed as a geometric vector. When θ_1_ and θ_2_ approach 90 degree, both vector *α* and vector *β* will be statistically uncorrelated with vector *η*. However, *η* could be a linear function of *α* and *β* when and only when the three vectors are in the same plane (Scenario 2). **(b)** The practice of uncorrelated-trait-based modeling depends on the convergence/divergence nature of the phenotype space. If the space is divergent, the number of sampled dimensions would increase infinitely with trait sampling. As a result, the sampled traits would each have unique dimensions and the trait space is logically splitting. On the contrary, the number of sampled dimensions would reach a plateau if the space is convergent. As a result, there would be a lot of shared dimensions among the sampled traits such that the trait space is logically fully interconnected. Hence, good practice of the trait modeling is more likely in a convergent space.

When the angle approaches 90 degree, the correlation will approach zero and thus the two vectors would be statistically uncorrelated. In the first scenario where *η* is nearly perpendicular to the plane defined by *α* and *β*, there is little power for α and β to model *η*. This is because neither *α* nor *β* can model the Z-dimension of *η*. In contrast, in the second scenario where *η* is in the same plane as *α* and *β, η* can be well described by a function of *α* and *β* although they are as uncorrelated as in the first scenario to *η*. Hence, capturing all dimensions of a high-dimensional trait is the key to successful modeling, while the two-dimensional measure of correlation alone does not tell much about a high-dimensional space.

In practice, a trait is a statistic sampled from a phenotype space. As shown in Fig. 3b, because a trait, say *η*, cannot be well represented by a single uncorrelated trait, say, *α*, sampling another trait in the phenotype space, say, *β*, would be required for modelling *η*. However, *β* may not be at exactly the same plane as *α* and *η*, so an extra trait, say, *γ*, that can represent the *β*-specific new dimension would be required for modelling *β*. Similarly, sampling an additional trait that can represent the *γ*-specific new dimension might be required for modelling *γ*. This process could continue to have a large number of traits sampled. If the space is convergent with a finite number of basic dimensions, there would be a situation where the basic dimensions are recurrently used in the sampled traits such that even seemingly unrelated traits are logically interconnected. On the contrary, if the space is divergent with infinite dimensions, the sampled trait space would most likely be splitting with no shared logics between uncorrelated traits.

We modelled all the remaining traits (405–20=385), respectively, using their uncorrelated traits. We achieved similar successes as in the 20 exemplar traits. Specifically, the variances explained by the linear models are overall comparable to *H* ^2^ of the dependent traits (Fig. 4a and Fig. S12). Because *H*^2^ also represents the repeatability of trait measuring, it may approximate the upper bound of the modeling performance. Hence, the space formed by the 405 traits appears to comprise virtually all logics required for modeling the available traits. This is a signature of convergent space where all objects can be logically interconnected with each other. In contrast, the performance of modeling the yeast traits using their uncorrelated gene expressions is consistently poor (Fig. S13), highlighting the logic stratification between the two spaces.

**Fig. 4.**
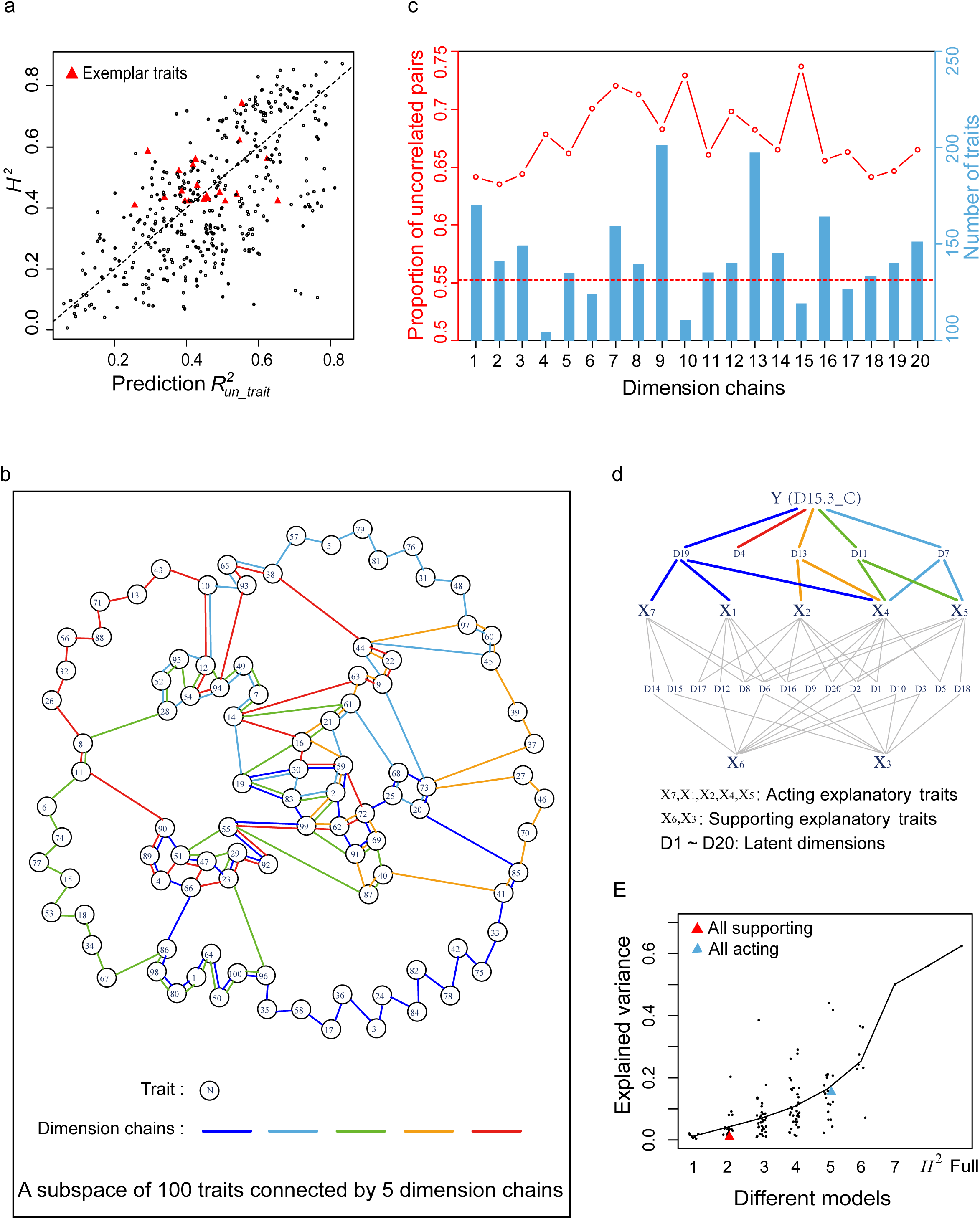
Analysis of dimension chains. **(a)** The prediction performance 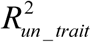 is overall comparable to *H*^2^ for the 405 yeast traits. **(b)** A subspace comprising 100 traits connected by five dimension chains. The 100 traits are selected because the absolute Pearson’s *R* between any two of them is less than a given value (0.65), and the five dimensions are arbitrarily chosen from the 20 latent dimensions. Traits sharing a latent dimension are connected by a “chain” of the given dimension. **(c)** The number of traits connected by each of the 20 latent dimension chains is shown, with the fraction of uncorrelated trait pairs in the chain provided. **(d)** A bipartite network showing how D15.3_C is connected with its top seven uncorrelated explanatory traits via the latent dimensions. One dimension (D4) of D15.3_C cannot be effectively modelled by the seven explanatory traits. There are five acting explanators (X_7_, X_1_, X_2_, X_4_, and X_5_) that share at least one dimension with D15.3_C and two supporting explanators (X_6_, and X_3_) that share no dimension with D15.3_C. Both types are important for the modeling of D15.3_C. **(e)** The variance explained by the nested models with only some (*n* = 1, 2… 7) of the top seven explanatory traits of D15.3_C remained. There are 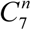 nested models examined for each *n* category. For *n* = 1, the variance explained by the models is equal to the square of the Pearson’s correlation coefficient of each explanatory trait with D15.3_C. The results are compared to *H*^2^ of D15.3_C as well as to the variance explained by the full model for D15.3_C.

The general success of the uncorrelated-trait-based modeling suggests long-distance logic chains prevailing the phenotype space. Since the logics underlying traits are by nature dimensions of the phenotype space, we analyzed the latent dimensions owned by each of the 405 traits, attempting to demonstrate the logic/dimension chain structure. According to the modeling results we had:

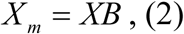

where *X* is the 815 by 405 phenotype matrix, in which each row represents a segregant and each column represents a trait, *B* is the 405 by 405 coefficient matrix, in which each column is the learned coefficient vector of a dependent trait and only uncorrelated traits of the focal dependent trait could have non-zero coefficients and *X* _*m*_ is the modelled portion of *X*. Using a deep-learning approach to reduce dimensions (Methods) we obtained

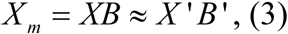

where *X*’ is an 815 by 20 matrix, in which each row represents a segregant and each column represents a latent dimension, *B*’ is a 20 by 405 coefficient matrix, in which each column is a coefficient vector of a trait. Approximately 85% variance of *X* _*m*_ can be explained by *X*’*B*’ (Methods). Hence, the 405 traits are each approximated by an additive function of 20 latent dimensions. The relatedness of two traits is well explained by the similarity of their latent dimension spectra in *B*’ (Pearson’s *R* = 0.76 for all trait pairs; Fig. S14a). Tow traits are connected via a latent dimension if the dimension is non-trivial for both of the traits under a stringent cutoff (Fig. S14b and c; Methods). This way, each latent dimension could form a “chain” to connect all the traits that own the dimension, and the whole trait space becomes fully interconnected via the 20 dimension chains (Fig. 4b and Table S1). On average a trait is passed through by seven dimension chains (Fig. S14d) and a dimension chain connects as many as 144 traits (Fig. S14e), most of which are uncorrelated (Fig. 4c). Up to 94% of the trait pairs are directly connected via at least one dimension chain and the rest require a third trait to bridge them (Fig. S14f). With these properties the dimension chains account for the linear dependence of uncorrelated traits in the phenotype space.

A close examination of the latent dimensions owned by each trait revealed additional insights. Again we use the trait D15.3_C as an example: Among the seven top explanatory traits five share at least one dimension with D15.3_C while two share no dimension with D15.3_C (Fig. 4d). Hence, the various dimensions of D15.3_C are modelled directly by acting explanatory traits, which, however, would introduce additional dimensions that require supporting explanatory traits to cancel them out. Notably, both the acting and supporting explanatory traits are important to the success of the modeling, as evidenced by testing the nested models that consider only some of the explanatory traits (Fig. 4e). This process is like what Fig. 3a shows: neither vector *α* nor vector *β* alone describes vector *η* well, but *α* and *β* together can fully describe *η* so long as they have all required dimensions.

## Discussion

In this study we demonstrate the interconnection of a phenotype space via long-distance dimension chains. These phenotypic dimension chains can nearly fully bridge even statistically uncorrelated traits, enabling precise inferences based on seemingly unrelated traits. Notably, focusing on correlated traits, as what conventional studies have done, would break such long-distance chains into pieces, resulting in “splitting” understandings of a phenotype space. Another major discovery is the dimension stratification between genotype space (broadly defined as genome plus transcriptome) and phenotype space, as evidenced by the lack of shared dimensions between traits and SNPs/expressions. In fact, we found uncorrelated gene expressions of the yeast are also linearly dependent (Fig. S15), suggesting similar interconnection within the expression space. Hence, the macroscopic layer and microscopic layer of the yeast each appear to be a self-explanatory linear space, with limited cross-space dimensions available (Fig. 5). While our analyses are based on yeast, we see no reason that the principles revealed are exception given that the expression space examined is composed of all ∼6,000 yeast genes and the phenotype space examined is composed of several hundred traits with all kinds of statistically uncorrelated pairs (Fig. S16).

**Fig. 5.**
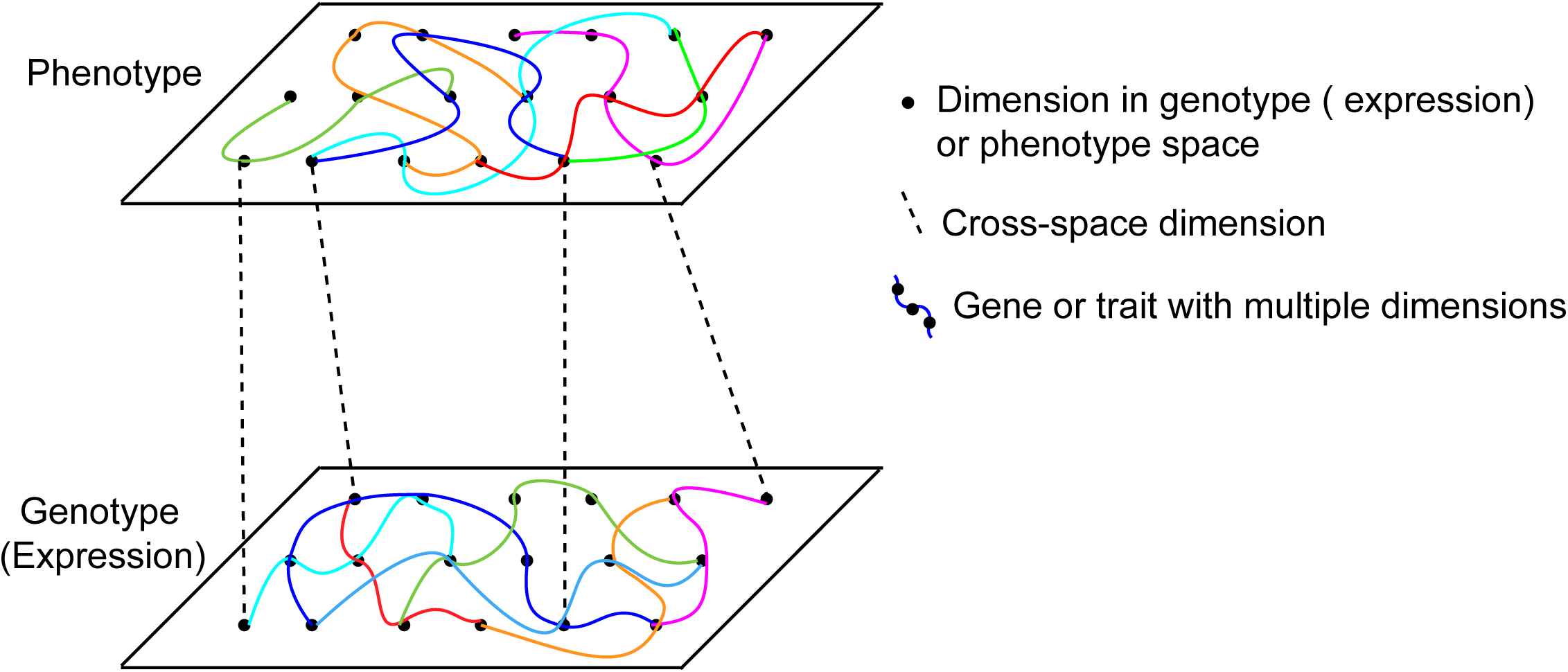
The dimension stratification between genotype (expression) space and phenotype space. In each space there is a set of dimensions recurrently used to represent various traits (or gene expressions), which underlies the success of within-space inferences. In contrast, between-space inferences are often difficult because of the limited cross-space dimensions. In yeast, cross-space dimensions can only explain *h*^2^, and the unexplained trait variance is due to the lack of appropriate dimensions.

Given the world we are living is phenotypic and what people care about is phenotype, the ability of making precise inferences across whole phenotype space calls for a revisit of the current genotype-centered views in biological research. More importantly, the logic stratification between genotype and phenotype predicts the unavoidable difficulty of the bottom-up genetic thinking in dealing with complex traits. Hence, developing phenotype-centered views may provide a valuable complement to current genetics(*18*). We conclude this article by outlining a few next challenges. First of all, it is unclear how the ideas presented would be adapted to multicellular organisms, since the phenotype space would be composed of various layers, ranging from cell, tissue, organ, to whole body, as well as various types, including morphology, physiology, behavior, and so on. It is possible that the basic dimensions of such a phenotype space are clustered according to the layers or types, which would affect the practice of making trait inferences in a sampled subspace. It would a fascinating field to explore how we could collect all the basic phenotypic dimensions of a complex organism such as humans. While the potential differences are stressed, we note that the total number (∼6,000) of genes of the yeast is comparable to that of most complex organisms (e.g., ∼25,000 genes for humans). It is possible that the basic dimensionality of phenotype space might also be of comparable scales for them. Second, the from-trait-to-trait models described in this study are explanatory. They are helpful for predicting a trait when the trait and its closely related traits are not accessible (e.g., predicting milk productivity from the morphology of a baby cow). However, it is unclear how they would help modify a trait. Further efforts are required to transform the explanatory models into causal ones, which would help the design of trait modifications within phenotype space (e.g., modifying body weight to avoid cardiovascular disease, or physical therapy such as massage). Third, the basic dimensions of a phenotype space are so far mathematical abstractions. It would be interesting to reveal their biological nature.

## Methods

### Data

We studied a panel of segregants of a yeast hybrid (*S. cerevisiae* strain BY x strain RM) generated by a previous study (Bloom et al, Nature, 2013)(*15*). A total of 1,008 segregants are available with genotype information, among which 815 were phenotyped (*16*). In brief, the yeast cells were first stained by two fluorescent dyes, one for staining cell wall and the other for staining nucleus. Then, the microscopic images of the stained cells were obtained by a high-throughput automated image capturing system IN Cell Analyzer 2200 (GE Healthcare). The images were analyzed by a computer software CalMorph, and a total of 405 quantitative traits related to the yeast cell morphology were obtained (*14*). The traits include the areas and circumferences, the elliptical approximation, brightness, thickness, axis length, neck width, neck position, bud position, axis ratio, cell size ratio, outline ratio, proportion of budded cells, proportion of small budded cells, segment distances between mother tip, bud tip, middle point of neck, center of mother and bud, nuclear gravity centers and nuclear brightest points and angles between segments in different cell stages and so on. The phenotyping was conducted twice for each of the segregants, and trait values were Z-score transformed. Among the 815 phenotyped segregants, 796 have RNA-seq-based gene expression profiles that were generated by a recent study (*17*). The file AllData.xlsx contains the details of all data used in this study.

### Estimation of heritability

#### Broad-sense heritability (H ^2^)

Following Bloom et al (Nature, 2013), a focal trait is modelled by linear mixed model (LMM) as

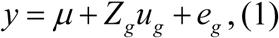

where *y* is the focal trait, *µ* is the population mean, *Z*_*g*_ is the design matrix indicating which segregant each replicate belongs to, *u*_*g*_ is a vector of random effect, and *e*_*g*_ is a vector of residuals. The variance of the foal trait is decomposed into genetic effect 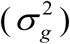 and environmental effect 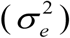. *H* ^2^ is then estimated as 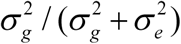. The ‘lmer’ function in R package ‘lme4’ is used. Standard error is estimated by Jackknife.

#### Narrow-sense heritability (h^2^)

Following Bloom et al (Nature, 2013), a focal trait is modelled by LMM as

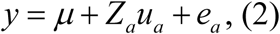

where *Z*_*a*_ is the identity matrix, *u*_*a*_ is a vector of random effect, 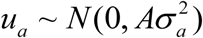, and *e*_*a*_ is a vector of residuals. The variance structure of the trait is formulated as

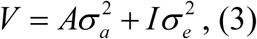

where A is estimated as the relatedness matrix between two segregants, which is calculated by the function ‘A.mat’ in R package ‘rrBLUP’, *I* is the identity matrix, 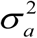 and 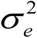 are additive variance and residual variance, respectively, which are estimated by R package ‘rrBLUP’. Then, *h*^2^ is calculated as 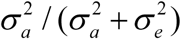. Standard error is estimated by Jackknife. When *h*^2^ is compared with the variance explained by quantitative trait loci (QTLs) or by uncorrelated SNPs, *y* refers to the average of two replicates. When *h*2 is compared with *H* ^2^ or the variance explained by uncorrelated traits, *y* refers to one randomly chosen replicate.

#### QTL heritability 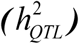

A focal trait with QTLs is modelled by multiple linear model as

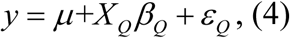

where *X* _*Q*_ is the QTL matrix, in which each row represents a segregant and each column represents a QTL, *β*_*Q*_ is the coefficient vector, and *ε*_*Q*_ is the residual vector. Then, 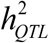 is estimated as the square of Pearson’s *R* between predicted and observed *y*. To obtain a robust estimation we conducted this modelling by machine learning. Specifically, each time the segregants are randomly divided into a training set composed of 715 segregants and a testing set of 100 segregants. The model is trained with 10-fold cross validation and L_2_ regularization, and then tested in the testing set. We repeated the process 20 times to obtain the mean and standard deviation. The function ‘cv.glmnet’ in R package ‘glmnet’ is used.

### QTL mapping and estimation of linkage blocks

We focused on 212 traits with *H*^2^ larger than 0.4. Following Bloom et al (Nature, 2013), the linkage between a SNP and a trait is estimated by LOD score defined as -*n*(ln(1-*R*^2^) / 2ln(10)), where *R* is Pearson’s correlation of them, and *n* (=815) is the number of segregants. An LOD score matrix for all SNP-trait pairs is obtained. To empirically estimate the statistical significance of LOD scores, we randomly shuffled the segregants to obtain a pseudo-LOD score matrix, and this process was repeated 1,000 times. The expected proportion of significant LOD scores is estimated under a series of cutoffs to derive the false discovery rate (FDR). We select the LOD score cutoff at which FDR is < 0.05 for QTL mapping. In the first round the most significant markers of the different chromosomes are selected; then, the residual trait value estimated by R package ‘qtl’ is used for the second round of QTL mapping. After two rounds of mapping, no significant signals remain. In the end, 210 of the 212 traits have at least one QTL mapped.

The Pearson’s *R*^*2*^ of every two SNPs in the segregant population is first calculated. For a given SNP we seek its nearest SNP that has a *R*^*2*^ < 0.1, and the segment between the two SNPs is regarded as a linkage block. After examining all SNPs, the mean and median size of the linkage blocks are 102,713bp and 106,025bp, respectively.

### Definition of exemplar traits

Among the 405 traits we selected a set of exemplars that are as uncorrelated as possible with each other. We first computed the Pearson’s *R* for all trait pairs. We adopted a heuristic method. Each time we deleted the trait that has the most number of correlated traits with absolute *R* > 0.35. We repeated the process until no trait pair has absolute *R* > 0.35, remaining 20 exemplar traits each with *H*^2^ > 0.4.

### Modelling a trait using its uncorrelated traits 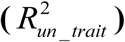

Two traits are called correlated if the square of Pearson’s *R* between them is larger than 0.02, which corresponds to *P* < 0.01 (T-test with Bonferroni correction). Based on this cutoff, the uncorrelated traits of a focal trait are obtained. A focal trait is then modelled as

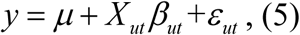

where *X* _*ut*_ is a matrix consisting of uncorrelated trait values, in which each row represents a segregant and each column represents an uncorrelated trait, *β*_*ut*_ is a coefficient vector, and *ε*_*ut*_ is a vector of residues. *β*_*ut*_ is estimated by machine learning. The learning process and the evaluation of prediction performance is the same as in the estimation of 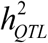, with the exception that a LASSO model (L_1_ regularization) instead of L_2_ regularization is used. Specifically, each time the segregants are randomly divided into a training set composed of 715 segregants and a testing set of 100 segregants. The model is trained with 10-fold cross validation and L_1_ regularization in the training set and then tested in the testing set. Prediction performance 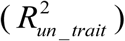 is measured as the square of Pearson’s *R* between predicted and observed *y*. We repeated the process 100 times for exemplar traits and 20 times for the other traits to obtain the mean and standard deviation for each of them. The function ‘cv.glmnet’ in R package ‘glmnet’ is used.

### Modelling a trait using its uncorrelated SNPs 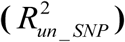

There are 11,623 SNPs included in the segregant population. A trait and a SNP are called correlated if the square of Pearson’s *R* between them is larger than 0.029, which corresponds to *P* < 0.01 (T-test with Bonferroni correction). At the same time, we also considered QTLs of a focal trait and their linked SNPs. A SNP is called linked with a QTL if they are significantly correlated in the yeast population under *P* < 0.01 (T-test with Bonferroni correction). Hence, the uncorrelated SNPs of a focal trait are those that are both not correlated with the trait and not linked with the trait’s QTLs. For each trait often ∼10% of the SNPs are excluded from the modelling. A focal trait is then modelled as

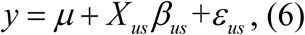

where *X* _*us*_ is the matrix consisting of uncorrelated SNPs, in which each row represents a segregant and each column represents an uncorrelated SNP, *β*_*us*_ is a coefficient vector, and *ε*_*us*_ is a vector of residues. The learning process and the evaluation of prediction performance are the same as in the modelling using uncorrelated traits. To be comparable with the modelling using uncorrelated traits, we required the number of uncorrelated SNPs used the same as the number of uncorrelated traits, by randomly selecting a subset of the uncorrelated SNPs. We repeated the process 100 times for each of the exemplar traits to obtain the mean and standard deviation of prediction performance 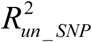. The function ‘cv.glmnet’ in R package ‘glmnet’ is used.

We also used LMM to model a focal trait with its total uncorrelated 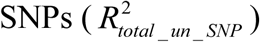. In our cases the number of uncorrelated SNPs appears too large to achieve a good performance in machine learning. The standard error of the modeling by LMM is estimated by Jackknife. The R package ‘rrBLUP’ is used.

### Modelling a trait using its uncorrelated gene expressions 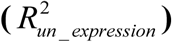

There are 5,720 genes included in an expression profile of the segregants. A trait and a gene expression are called correlated if the absolute Pearson’s *R* between them is larger than 0.168, which corresponds to *P* < 0.01 (T-test with Bonferroni correction). A focal trait is then modelled as

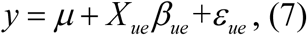

where *X* _*ue*_ is the matrix consisting of uncorrelated gene expressions, in which each row represents a segregant and each column represents an uncorrelated gene expression, *β*_*ue*_ is a coefficient vector, and *β*_*ue*_ is a vector of residues. The learning process and the evaluation of prediction performance are the same as in the modelling using uncorrelated traits. To be comparable with the modelling using uncorrelated traits, we required the number of uncorrelated gene expressions used the same as the number of uncorrelated traits used, by randomly selecting a subset of the uncorrelated gene expressions of a focal trait. We repeated the learning process 20 times for each of the exemplar traits to obtain the mean and standard deviation of prediction performance 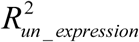. The function ‘cv.glmnet’ in R package ‘glmnet’ is used. We also used the same machine learning pipeline to model a trait using its total uncorrelated gene expressions 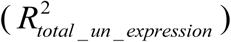, or using the whole gene expression profile 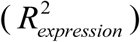.

### Modelling a gene’s expression using its uncorrelated gene expressions 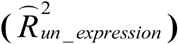

Two genes’ expressions are called correlated if the absolute Pearson’s *R* between them is larger than 0.150, which corresponds to *P* < 0.01 (T-test with Bonferroni correction). A focal gene’s expression is then modelled as

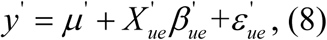

where y’ is the focal gene’s expression, 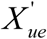 is the matrix consisting of uncorrelated gene expressions, 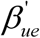 in which each row represents a segregant and 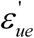 each column represents an uncorrelated gene expression, is a coefficient vector, and is a vector of residues. The learning process and the evaluation of prediction performance are the same as the modelling of traits. The function ‘cv.glmnet’ in R package ‘glmnet’ is used.

### Decomposition of latent dimensions of the phenotype space

There should be a set of latent dimensions for the 405 yeast traits. To uncover them, we first combined the learned *β*_*ut*_ in formula (5) for each of the 405 traits to form a matrix B with missing items assigned zero. We used *X* to represent the matrix consisting of the 405 traits, in which each row represents a segregant and each column represents a trait, and *X* _*m*_ to represent the modeled component of *X*. Hence, we had

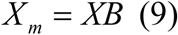

Let the matrix consisting of latent dimensions be *X*’, and the coefficients of traits on the latent dimensions form a new coefficient matrix *B*’, the Formula (9) can be rewritten as

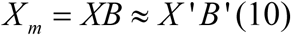

We solved (10) by autoencoder neural network, a routine deep-learning method. Specifically, we trained an autoencoder with both the input and output layers being *X* _*m*_, one hidden layer of 20 nodes being *X*’, and L_2_ regularization and sparsity regularization being used. The decoder transfer function was set as a linear function to form *B*’. The model explained nearly 85% variance of *X* _*m*_ when the hyper-parameter of L_2_ regularization is equal to 0.0001 and the sparsity proportion parameter equal to 0.5. As a result, *X*’ is an 815 by 20 matrix, in which each row represents a segregant and each column represents a latent dimension, and *B*’ is the decoder weight matrix. The MATLAB (2018b) function ‘trainAutoencoder’ is used.

In *X*’ the 20 latent dimensions are of comparable scales. Hence, the items in *B*’ weight the contribution of each latent dimension to a trait. To simplify the data structure in *B*’ we silenced the items by setting those smaller than 0.32 as zero. Under this cutoff, about two thirds of the items in *B*’ are silenced with 80% of the variance in *X*’*B*’ remained. Accordingly, a bipartite network connecting traits and latent dimensions is built using the R package ‘igraph’, and a dimension chain is to connect all the traits that have a link to a specific latent dimension in the bipartite network.

In principle, the latent dimensions could also be decomposed by conventional principal component analysis (PCA). We conducted PCA and found the top 20 PCs also explain ∼85% variance of *X* _*m*_. However, the explained variance of the top 20 PCs shows a much skewer distribution than that of the 20 latent dimensions decomposed by ‘autoEncoder’ (Fig. S17). As a consequence, the items in *B*’ would tend to be incomparable if PCs were used in *X*’, which would complicate the analysis of dimension chains. We therefore chose the latent dimensions decomposed by ‘autoEncoder’ instead of those by PCA. Notably, the choice affects only how the specific data (dimension chains) will be displayed rather than the main results.

## Figure legends

**Fig. S1.**
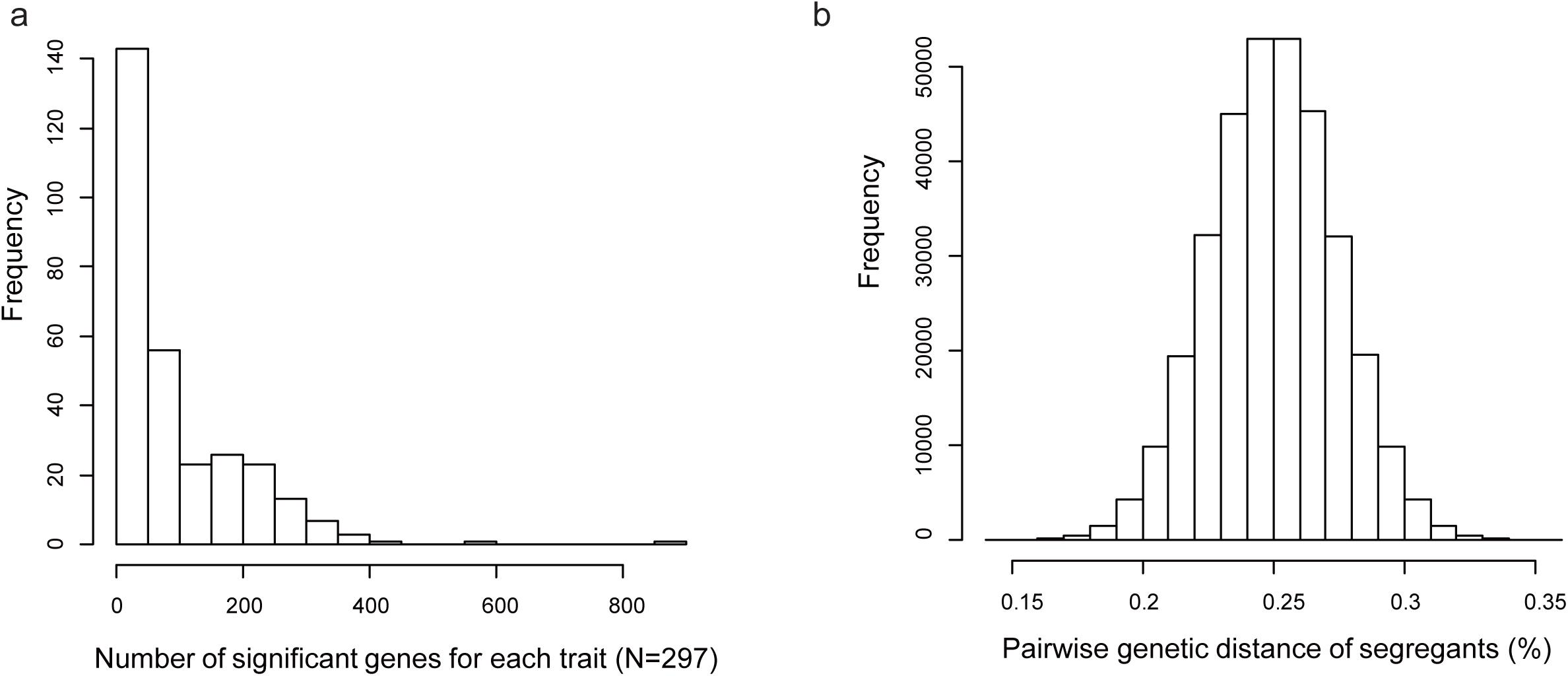
**(a) The frequency distribution of the number of genes that can significantly affect a morphological trait according to Ohya et al (PNAS, 2005)**. Among the 405 yeast traits examined in this study, 297 are suitable for the assessment (Threshold parameter = 0.3 in the Shapiro-Wilk test for normal distribution of wild-type trait values). A total of ∼2,700 non-redundant non-essential yeast genes are involved in regulating at least one of the 297 traits. **(b) The frequency distribution of pairwise genetic distance of the segregants**.

**Fig. S2.**
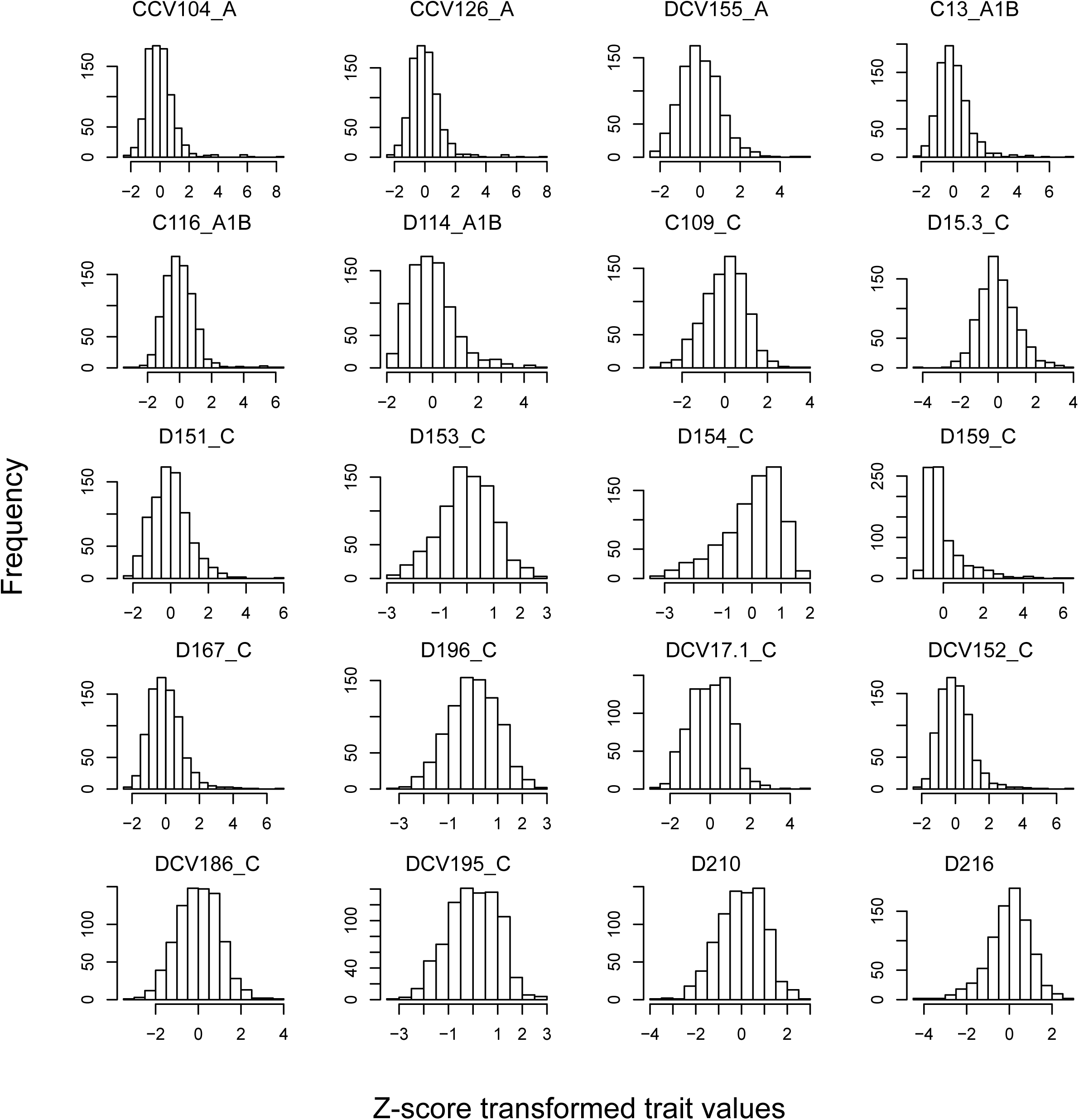
The bell-shape frequency distribution of the Z-score transformed trait values in the segregant population for each of the 20 exemplar traits.

**Fig. S3.**
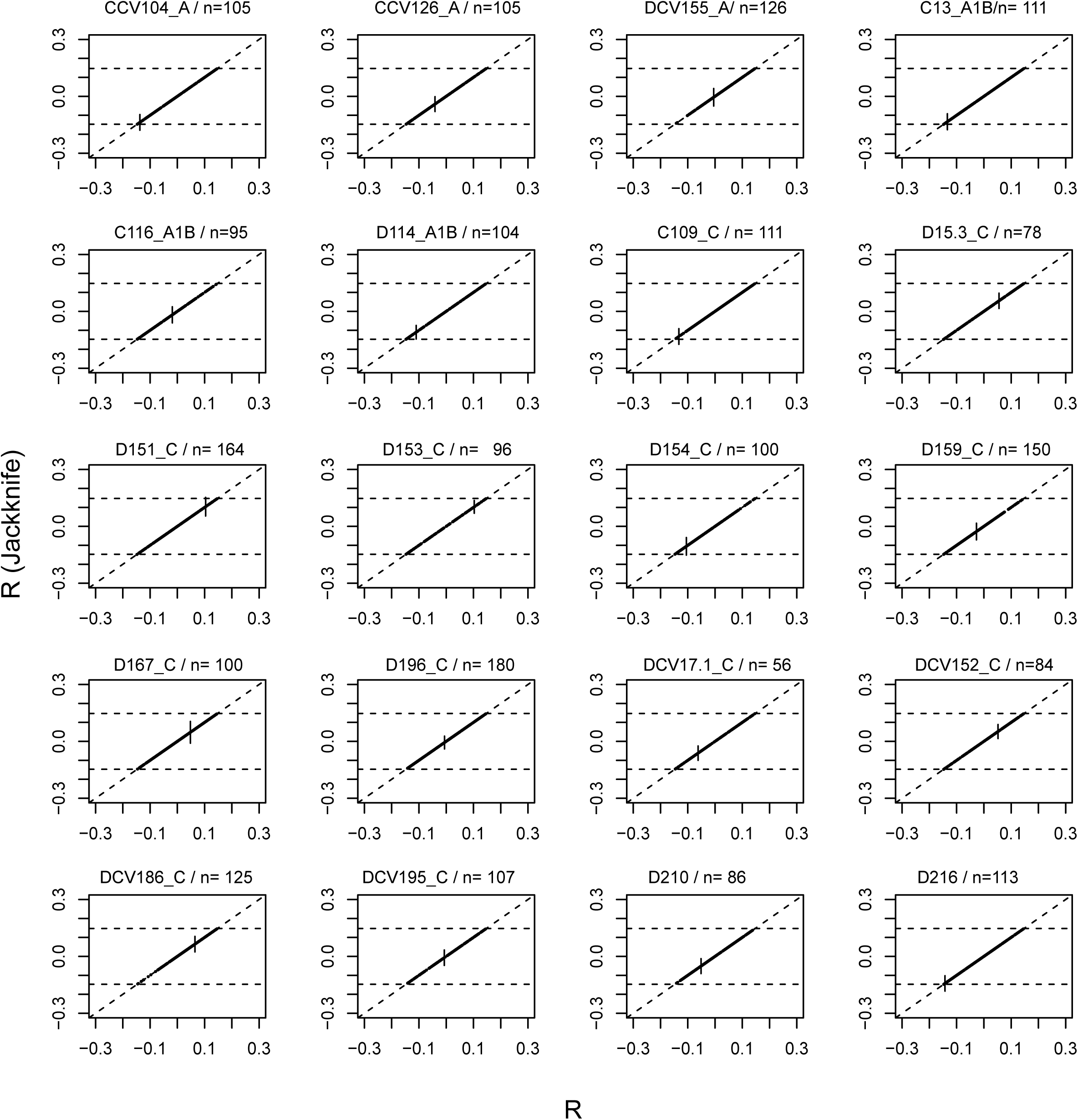
Robustness of the definition of uncorrelated traits of the 20 exemplar traits estimated by Jackknife method. Each panel represents an exemplar trait, with the number of uncorrelated traits (n) shown. The x-axis shows Pearson’s R between an exemplar trait and its uncorrelated traits in the segregant population. The y-axis shows the Pearson’s R estimated by Jackknife, with error bar showing the standard error. Horizontal lines in each panel show the cutoff used in the study to define an uncorrelated trait, and slant line shows y=x.

**Fig. S4.**
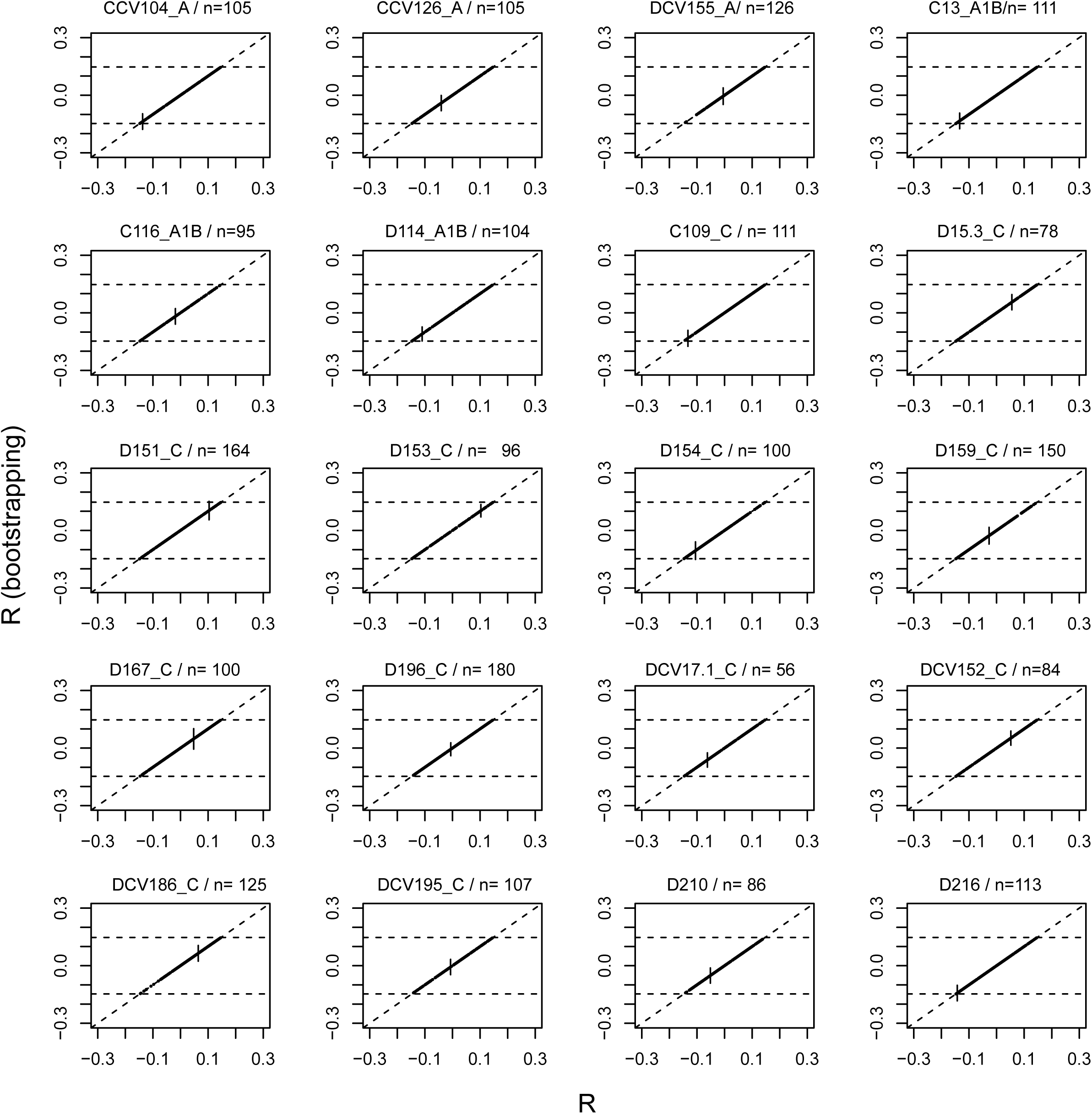
Robustness of the definition of uncorrelated traits of the 20 exemplar traits estimated by bootstrapping method. Same as Fig. S3 except that bootstrapping method is used to gauge the robustness. For each exemplar trait 1,000 bootstrapped samples are analyzed, and error bar shows the standard deviation.

**Fig. S5.**
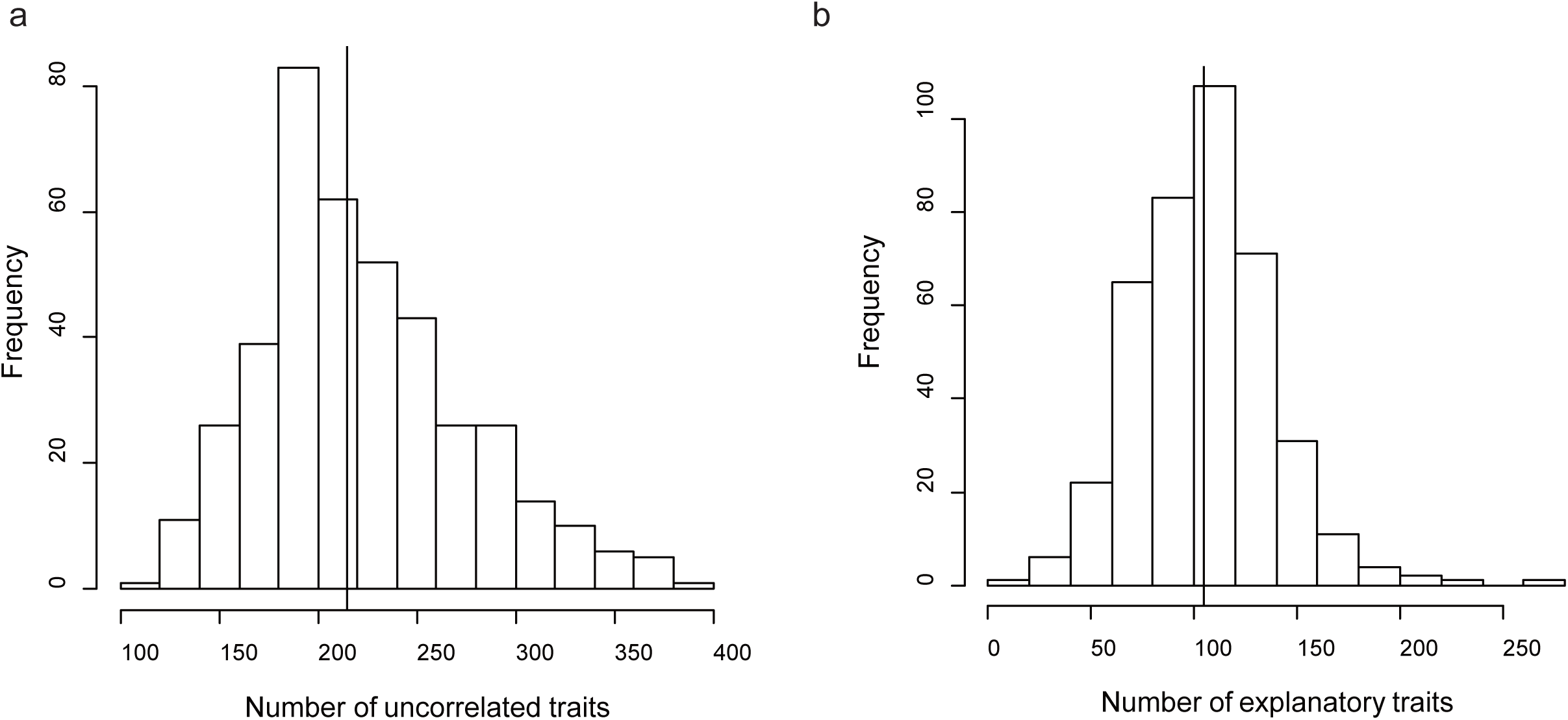
**(a) The frequency distribution of the number of uncorrelated traits of the 405 traits**. The vertical line shows the median. **(b) The frequency distribution of the number of explanatory traits for the 405 traits**. Explanatory traits here refer to those uncorrelated traits with non-zero coefficients in the linear function. The vertical line shows the median.

**Fig. S6.**
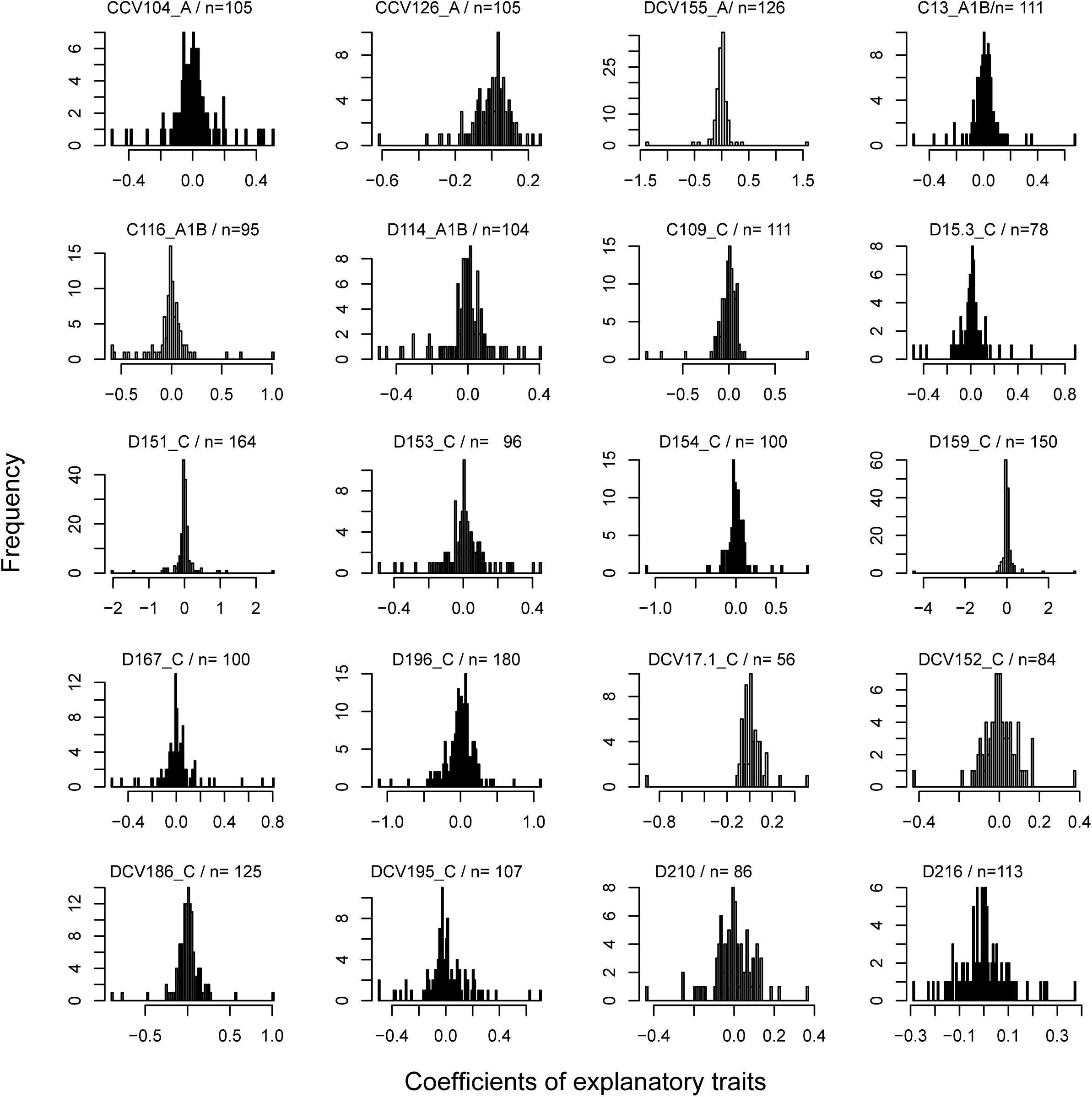
The frequency distribution of the coefficients of explanatory traits in the learned model of each exemplar traits. Each panel shows one exemplar trait, with the number of uncorrelated traits (n) shown.

**Fig. S7.**
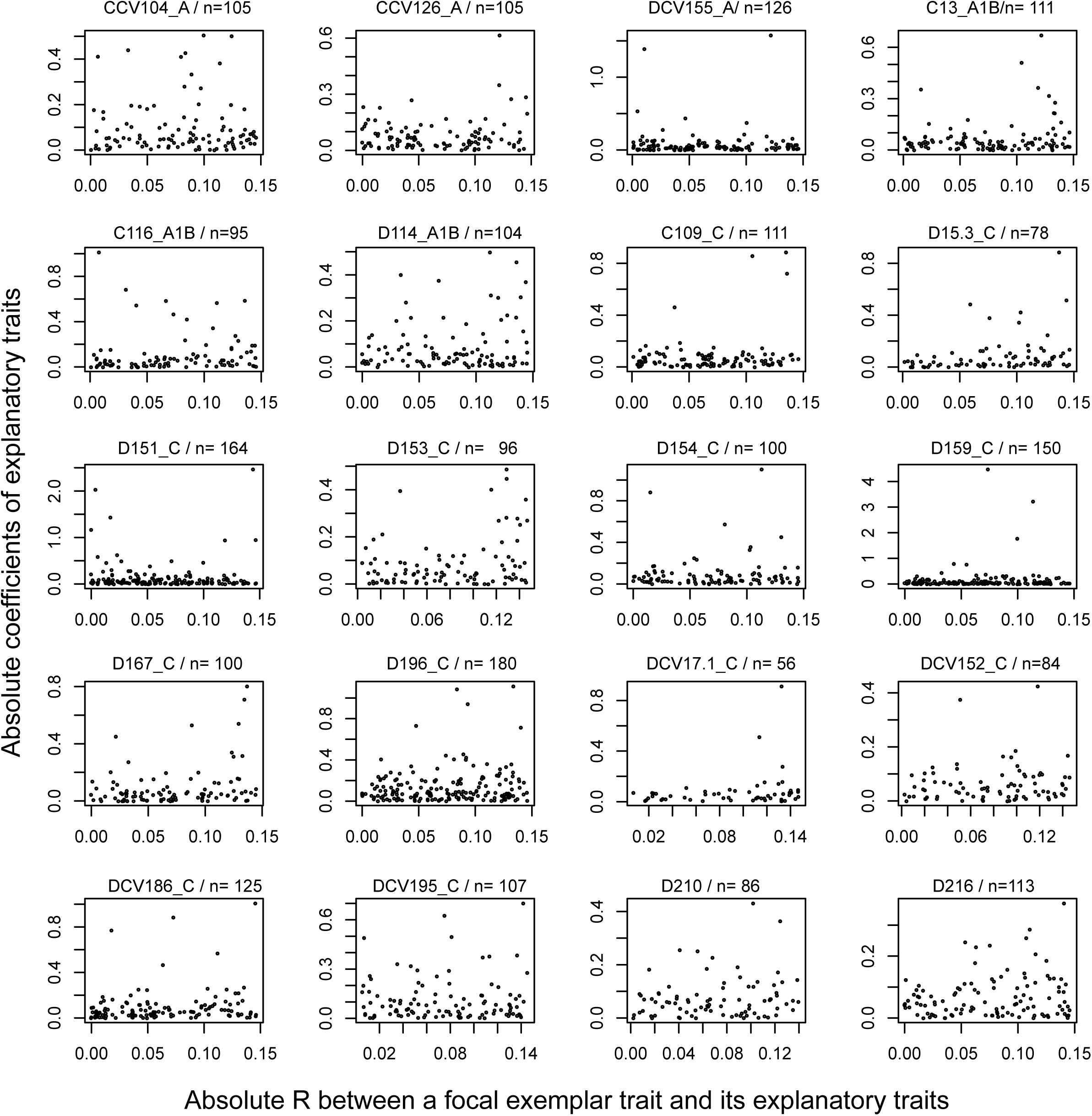
No apparent relationship between R (x-axis) and coefficients (y-axis) of the uncorrelated traits. Each panel shows one exemplar trait, with the number of uncorrelated traits (n) shown.

**Fig. S8.**
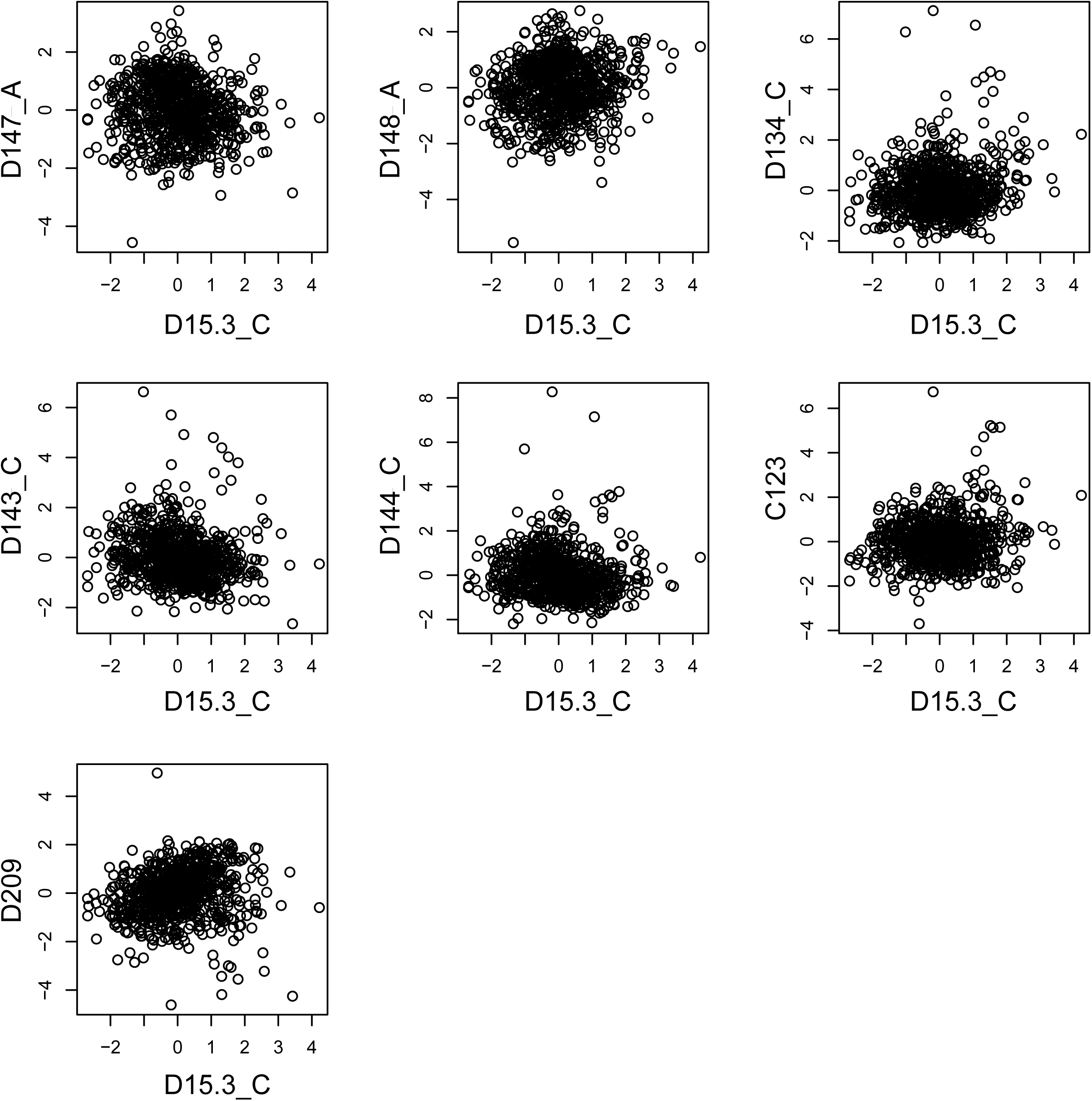
The scatter plot of the trait values of D15.3_C versus each of the top seven explanatory traits. No apparent outliers or data structure bias explain the uncorrelation between D15.3_C and the explanatory traits.

**Fig. S9.**
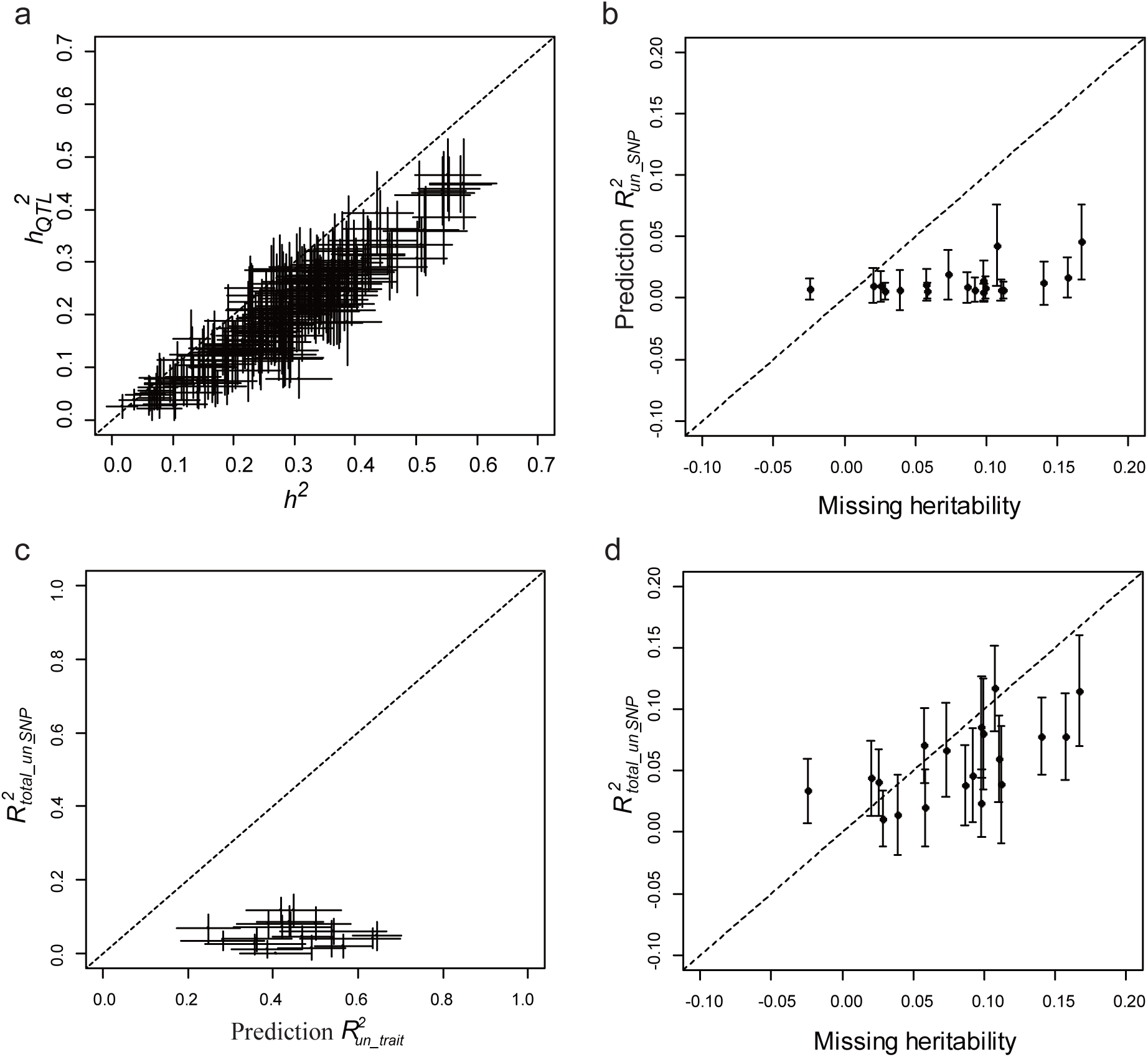
Much less trait variance explained by SNPs than by uncorrelated traits. **(a)** The relationship between 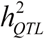 (heritability estimated by QTLs) and *h*^2^ (narrow-sense heritability) for 210 traits with at least one QTL mapped. **(b)** The relationship between 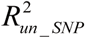 and missing heritability 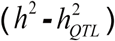 of the 20 exemplar traits. Here 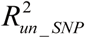 refers to trait variance explained by uncorrelated SNPs in the modeling, where the number of uncorrelated SNPs used is the same as that of uncorrelated traits used for modeling the exemplar traits. **(c)** The relationship between 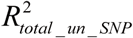 and 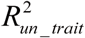 for each of the 20 exemplar traits. Here 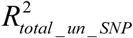 refers to trait variance explained by the total uncorrelated SNPs of a focal trait. **(d)** 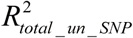 explains roughly the missing heritability 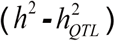 of the exemplar traits. The error bars of 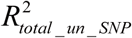 are standard error estimated by Jackknife. The error bars of 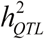 are standard deviation estimated by 20 repeats of learning. Slant dashed line shows y=x.

**Fig. S10.**
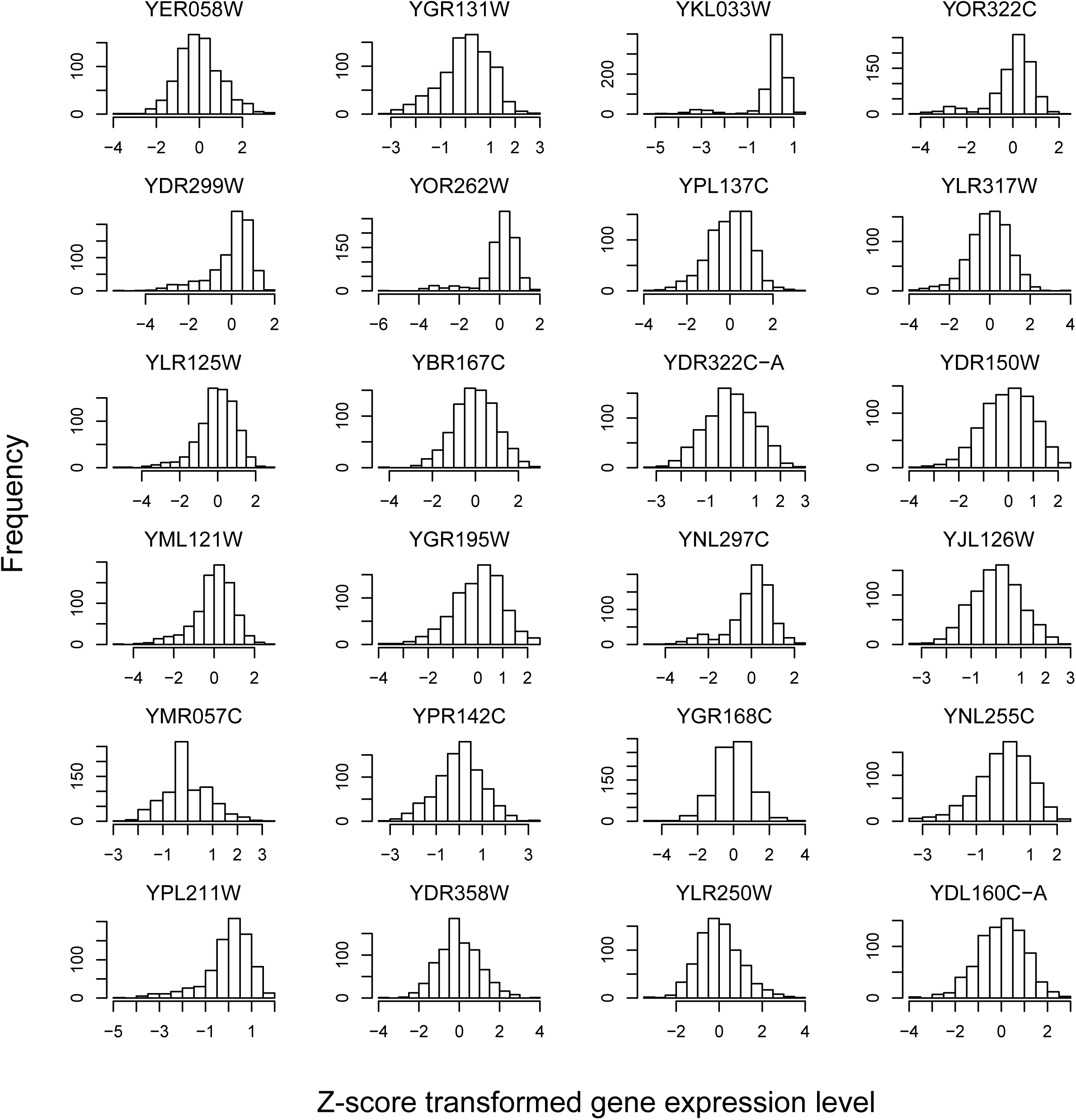
The bell-shape frequency distribution of Z-score transformed gene expression levels in the segregant population for 24 randomly selected genes.

**Fig. S11.**
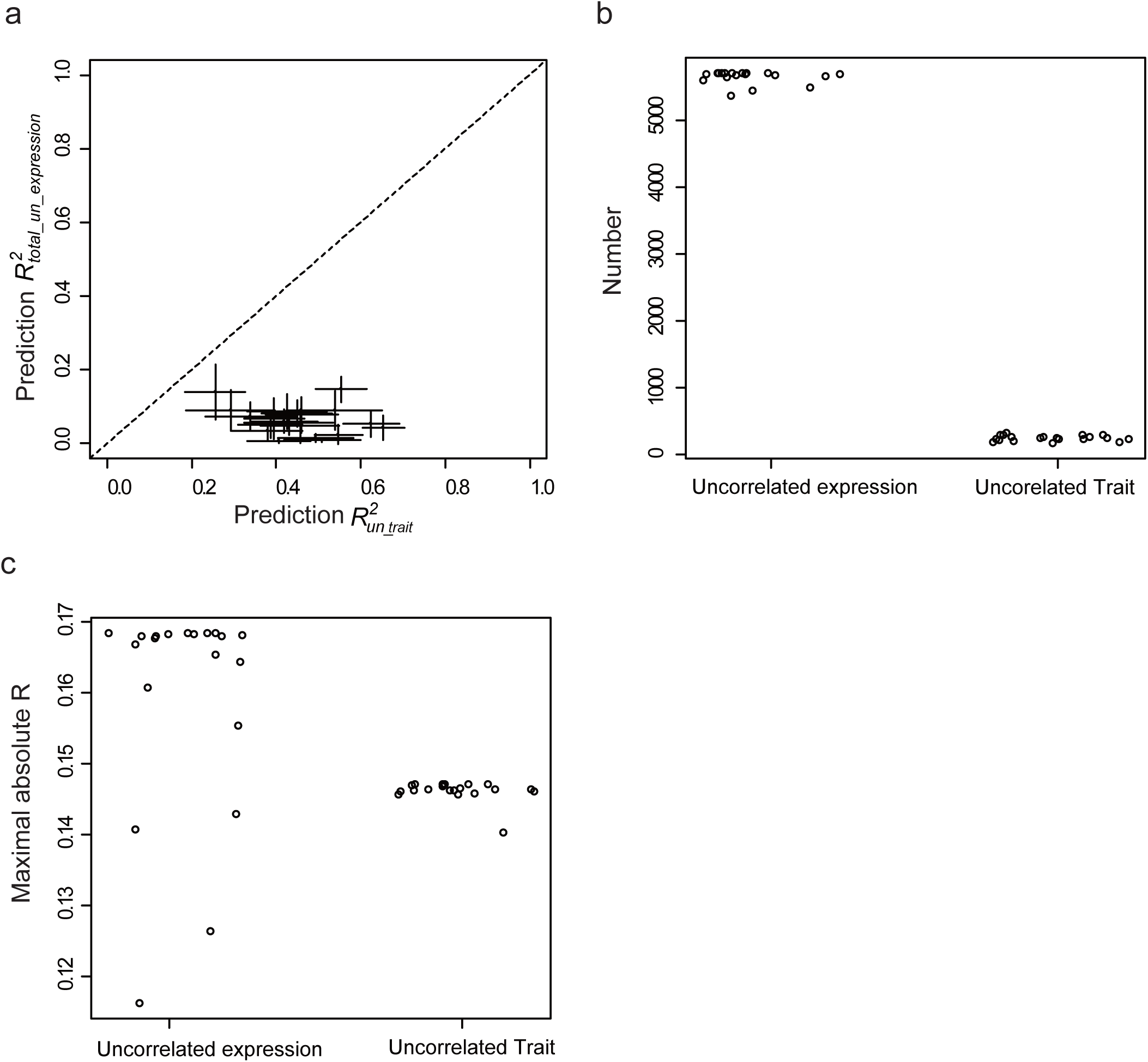
Much less trait variance explained by gene expressions than by uncorrelated traits. **(a)** The relationship between 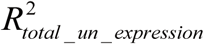 and 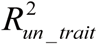. Here 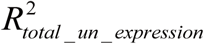 refers to trait variance explained by the total uncorrelated gene expressions of a focal exemplar trait. Error bars of 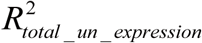 are standard deviation estimated by 20 repeats of learning. **(b)** Comparison of the number of uncorrelated gene expressions and the number of uncorrelated traits of the 20 exemplar traits. **(c)** Comparison of the maximal absolute Pearson’s R between a focal trait and its uncorrelated traits and the maximal absolute Pearson’s R between a focal trait and its uncorrelated gene expressions. The 20 exemplar traits are showed.

**Fig. S12.**
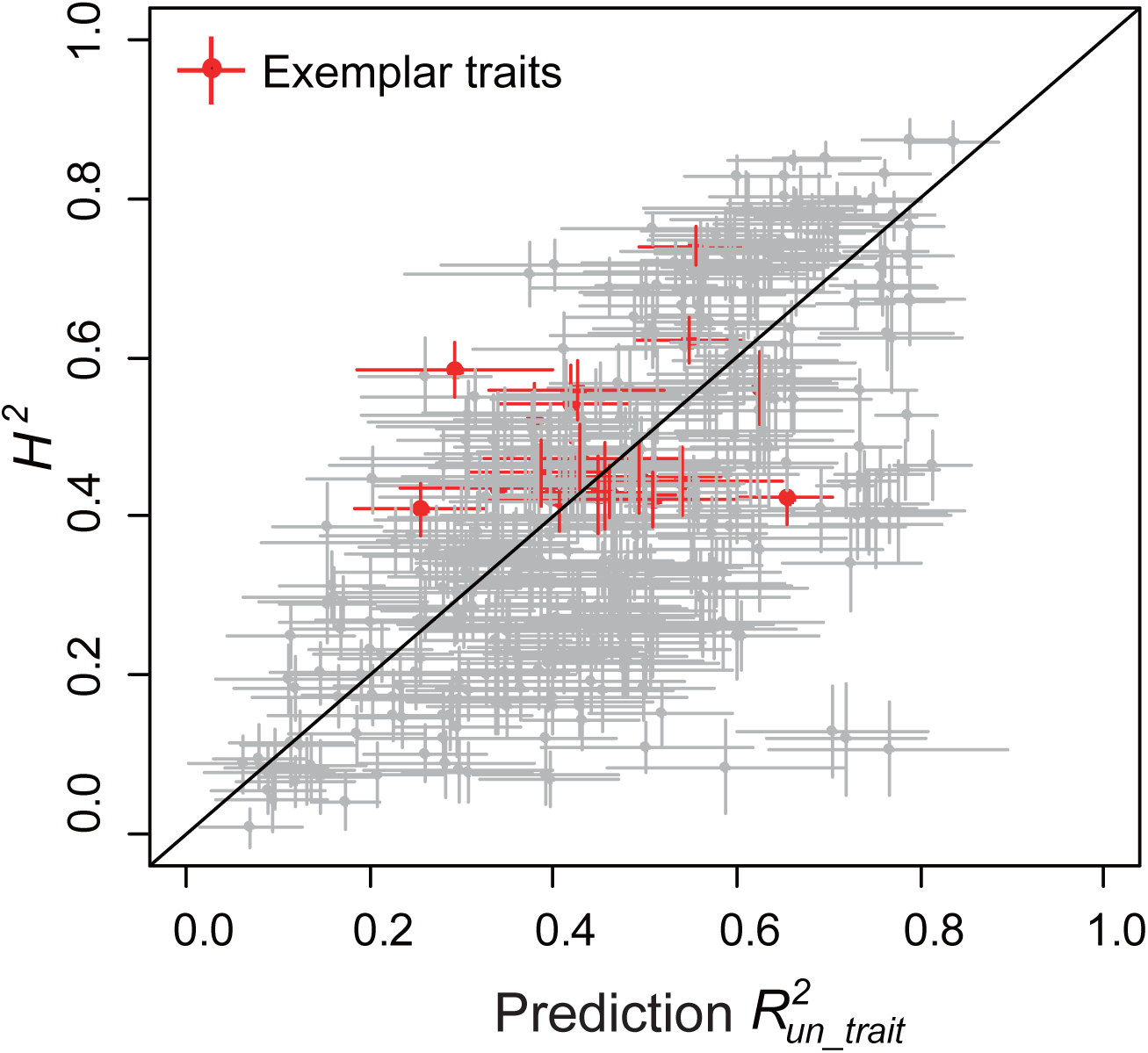
The prediction performance 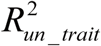 is overall comparable to *H*^2^ for the 405 yeast traits. Same as Fig. 4A except that error bars of the estimations are included. Error bars of *H*^2^ are standard errors estimated by Jackknife and error bars of 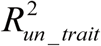 are standard deviations estimated by 20 repeats of learning.

**Fig. S13.**
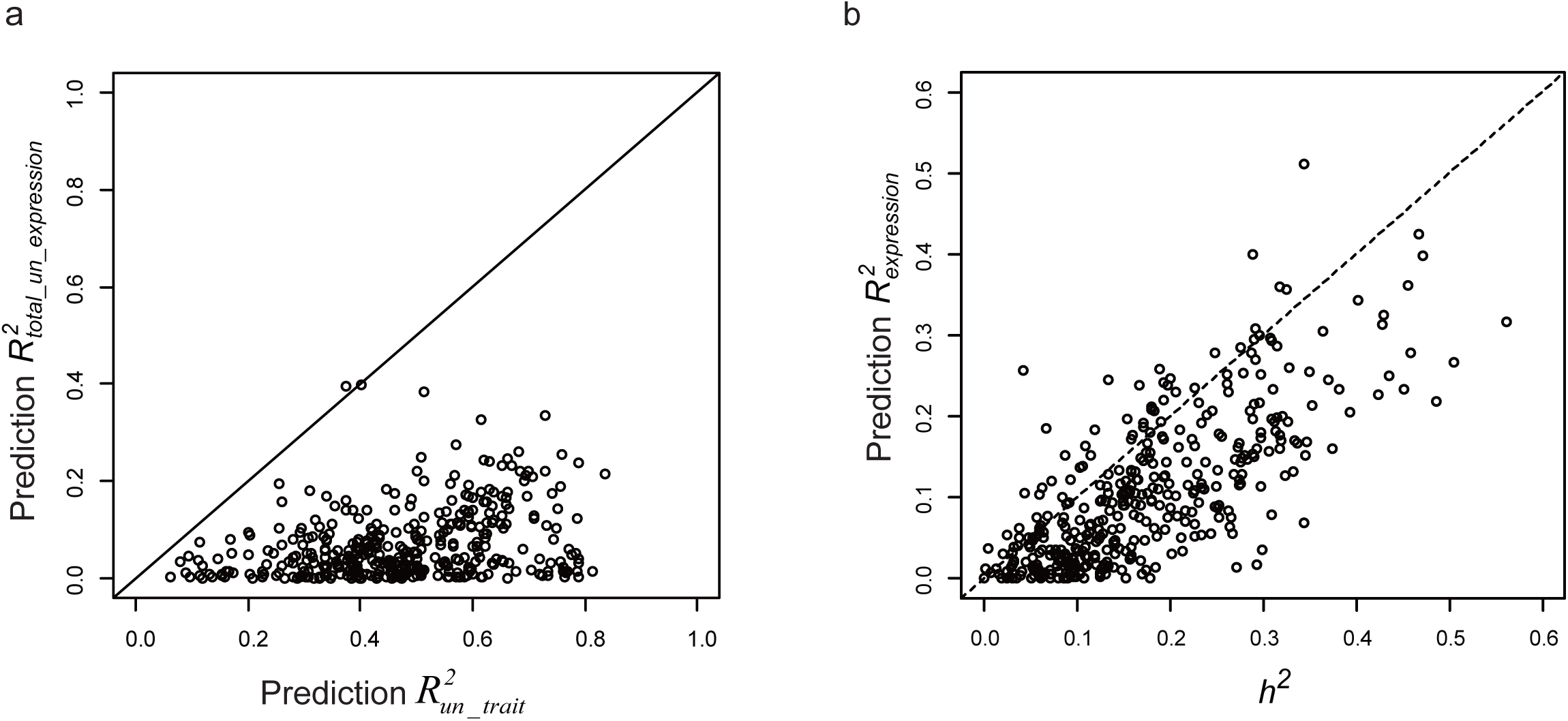
(a) 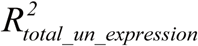 is consistently poorer than 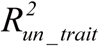 for the 405 traits. (b) 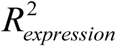 explains roughly the narrow-sense heritability (*h*^2^) of the 405 yeast traits. Here 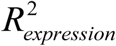 refers to trait variance explained by the whole gene expression profile.

**Fig. S14.**
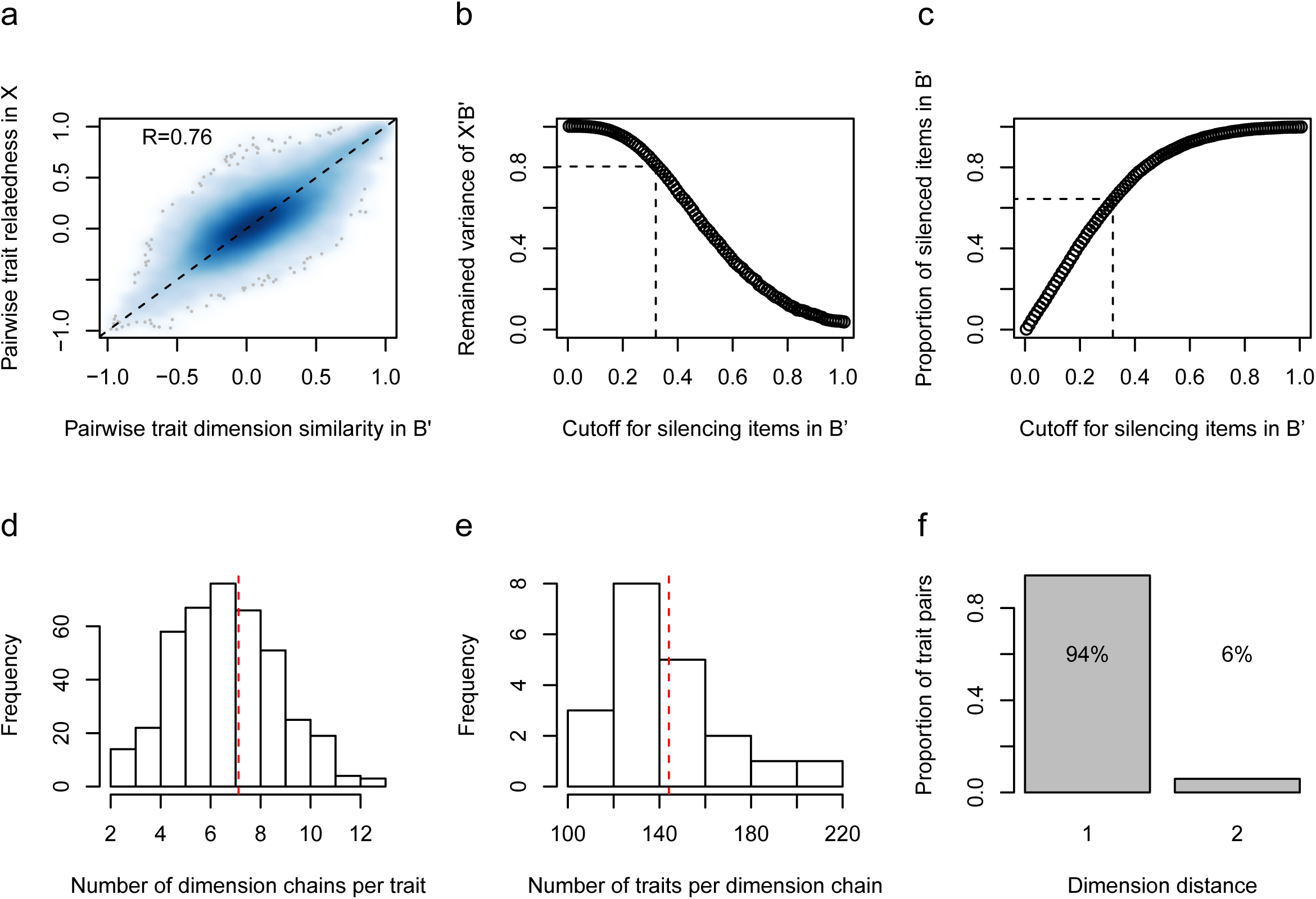
Features of latent dimension chains. **(a)** The relatedness of two traits (y-axis) is well explained by the similarity of their latent dimension spectra in *B*’ (x-axis). R refers to Pearson’s correlation coefficient. (**b)** The remained variance of *X*’*B*’ with items in *B*’ silenced under a series of cutoffs. Dashed lines shows 80% of the variance remained under the cutoff 0.32, which is used to defined non-trivial dimensions of a trait. **(c)** The proportion of silenced items in *B*’ under a series of cutoffs. Dashed line shows about two thirds of the items silenced under the cutoff 0.32. **(d)** The frequency distribution of the number of dimension chains passing a trait. The dashed line shows the median. **(e)** The frequency distribution of the number of traits each dimension chain connects. The dashed line shows the median. **(f)** The dimension distance between two traits. The distance is equal to 1 if two traits share at least one dimension. The distance is equal to n if two traits are indirectly connected by n dimension chains through n-1 other traits. All trait pairs have dimension distance no larger than 2.

**Fig. S15.**
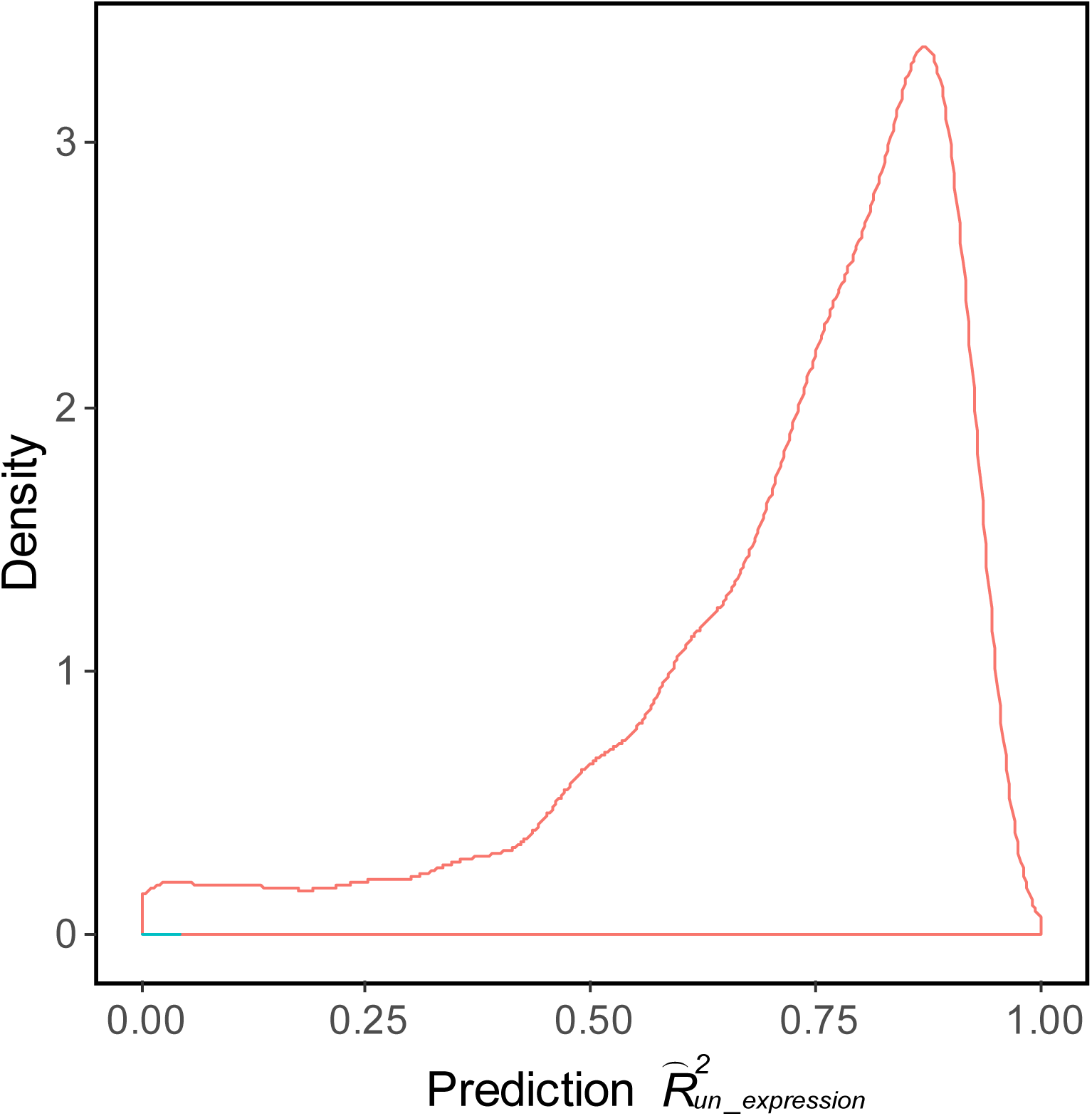
The density distribution of prediction performance of gene expression by uncorrelated genes expressions 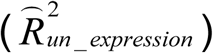 with a median of 0.77.

**Fig. S16.**
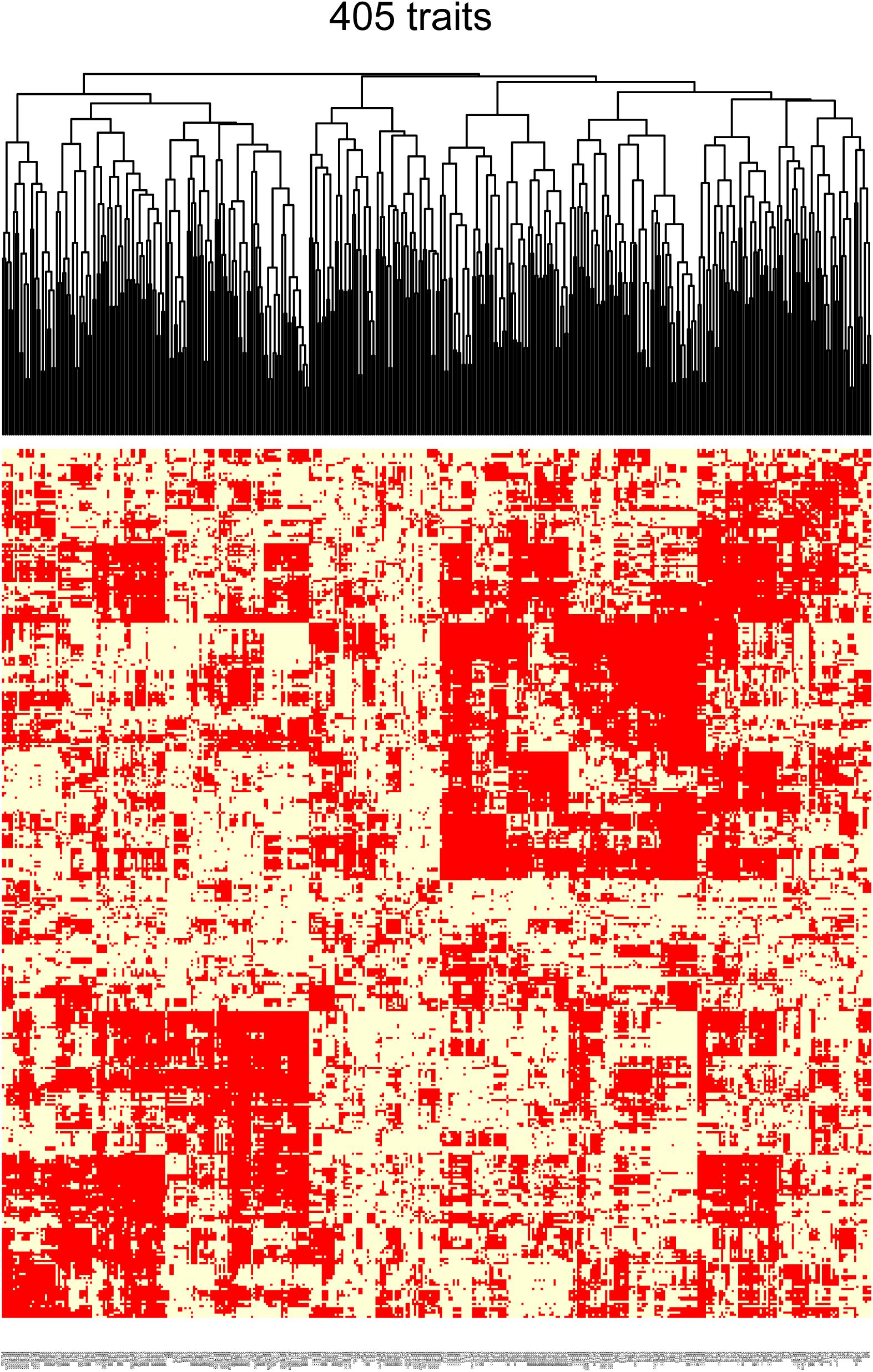
The yeast trait space examined in this study is composed of all kinds of uncorrelated trait pairs. The heatmap shows a highly heterogeneous distribution of uncorrelated versus correlated trait pairs. Uncorrelated trait pairs are labeled red and correlated trait pairs are labeled yellow. Traits are clustered based on the correlation profile similarity. The composition of uncorrelated traits varies substantially among the 405 yeast traits.

**Fig. S17.**
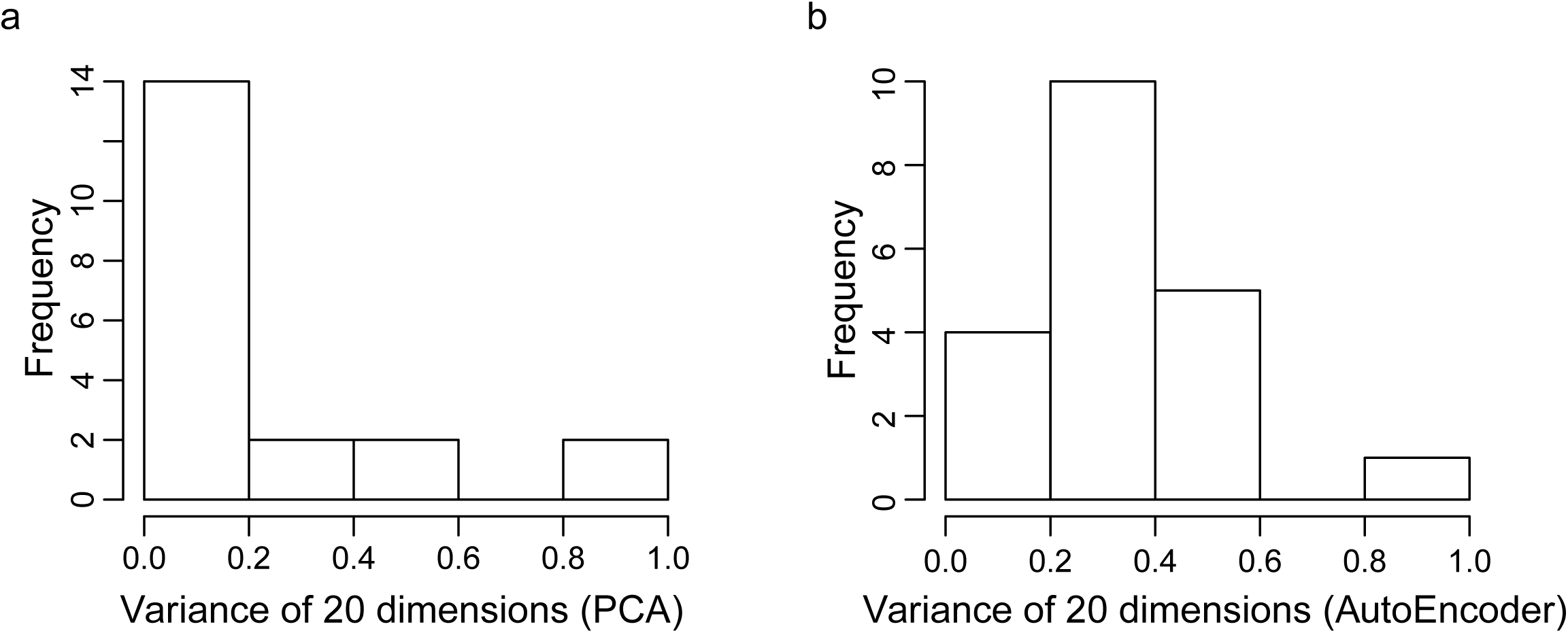
A and B are the distributions of variances of 20 latent dimensions defined by PCA and ‘autoEncoder’ methods, respectively. The minmax normalization is applied to both x-axes for comparison.

